# Classifyber, a robust streamline-based linear classifier for white matter bundle segmentation

**DOI:** 10.1101/2020.02.10.942714

**Authors:** Giulia Bertò, Daniel Bullock, Pietro Astolfi, Soichi Hayashi, Luca Zigiotto, Luciano Annicchiarico, Francesco Corsini, Alessandro De Benedictis, Silvio Sarubbo, Franco Pestilli, Paolo Avesani, Emanuele Olivetti

**Affiliations:** NeuroInformatics Laboratory (NILab), Bruno Kessler Foundation (FBK), Trento, Italy; Center for Mind and Brain Sciences (CIMeC), University of Trento, Italy; Department of Psychological and Brain Sciences, Indiana University, Bloomington, USA; PAVIS, Italian Institute of Technology (IIT), Genova, Italy; Neurosurgery Unit, Department of Neuroscience and Neurorehabilitation, Bambino Gesù Children’s Hospital, IRCCS, Roma, Italy; Division of Neurosurgery, Structural and Functional Connectivity Lab, S. Chiara Hospital, Trento, Italy

**Keywords:** white matter bundle segmentation, supervised learning, linear classification, diffusion Magnetic Resonance Imaging (dMRI)

## Abstract

Virtual delineation of white matter bundles in the human brain is of paramount importance for multiple applications, such as pre-surgical planning and connectomics. A substantial body of literature is related to methods that automatically segment bundles from diffusion Magnetic Resonance Imaging (dMRI) data indirectly, by exploiting either the idea of connectivity between regions or the geometry of fiber paths obtained with tractography techniques, or, directly, through the information in volumetric data. Despite the remarkable improvement in automatic segmentation methods over the years, their segmentation quality is not yet satisfactory, especially when dealing with datasets with very diverse characteristics, such as different tracking methods, bundle sizes or data quality. In this work, we propose a novel, supervised streamline-based segmentation method, called Classifyber, which combines information from atlases, connectivity patterns, and the geometry of fiber paths into a simple linear model. With a wide range of experiments on multiple datasets that span from research to clinical domains, we show that Classifyber substantially improves the quality of segmentation as compared to other state-of-the-art methods and, more importantly, that it is robust across very diverse settings. We provide an implementation of the proposed method as open source code, as well as web service.

## 1. Introduction

Accurate delineation of anatomical structures in the human brain is essential to numerous scientific disciplines. In particular, white matter bundle segmentation can provide information to multiple applications, e.g. the characterization of neurodevelopmental disorders, pre-surgical planning, or connectomic studies (Yeatman et al. (2012); O’Donnell et al. (2017); Yeh et al. (2018)).

In the last decade, several automatic methods for white matter bundle segmentation have been developed to mimic the manual segmentation done by expert neuroanatomists (Catani et al. (2002); Mori et al. (2005); Wakana et al. (2007)), which is very time consuming and difficult to reproduce. Automatic methods can be divided into three main groups: (i) Connectivity-based, (ii) Streamline-based, and (iii) Direct.

*Connectivity-based methods* aim to extract bundles by filtering the entire set of streamlines with inclusion/exclusion Regions of Interest (ROIs) that the bundle is assumed to pass / not to pass through (Oishi et al. (2008); Zhang et al. (2010); Yeatman et al. (2012); Wassermann et al. (2016)). These ROIs frequently come from atlases that have to be registered into the individual subject space. A significant drawback to this approach is that the segmentation is inherently limited by the anatomical variability of the subject, by the quality of the atlas, and by the process of registration.

*Streamline-based methods* group together streamlines according to some similarity measure. *Unsupervised* streamline-based methods, such as those in Brun et al. (2004); Maddah et al. (2005); O’Donnell and Westin (2007); Guevara et al. (2012); Tunç et al. (2014); Siless et al. (2016); Zhang et al. (2018), perform whole brain segmentation through clustering, without prior knowledge about the anatomy of the bundles and without leveraging examples of expert-made segmented bundles, limiting the quality of segmentation. In contrast, *supervised* streamline-based methods require one or more examples of the bundle to learn from, in order to segment such bundle in the target subject, such as those in Mayer et al. (2011); Olivetti and Avesani (2011); Vercruysse et al. (2014); Yoo et al. (2015); Labra et al. (2016); Garyfallidis et al. (2018) and Sharmin et al. (2018). It has been shown that streamline-based methods like those presented in Garyfallidis et al. (2018), referred to as RecoBundles, and in Sharmin et al. (2018), referred to as LAP, outperform connectivity based methods in terms of quality of segmented bundles.

*Direct methods* are voxel-based methods that segment bundles directly from diffusion images without the need for streamlines, see Wasserthal et al. (2018a) for a brief review. In contrast to the limited quality of segmentation reached by these methods, a recent direct method proposed in Wasserthal et al. (2018a) presented evidence of remarkably better segmentation quality in comparison with a large selection of other segmentation methods, including connectivity-based and streamline-based methods. This method, called TractSeg, is based on convolutional neural networks (Ronneberger et al. (2015)) and has set the new standard in terms of quality of bundle segmentation.

Despite the remarkable improvement in automatic segmentation methods over the years, the resulting bundles can be unsatisfactory. The quality of segmentation may be strongly affected by some properties of the bundles, for example by their size; by the tractography technique, e.g. probabilistic or deterministic tracking algorithm; or by the data quality, e.g. research (high-resolution) or clinical quality, see Figure 1 for some examples.

**Figure 1:**
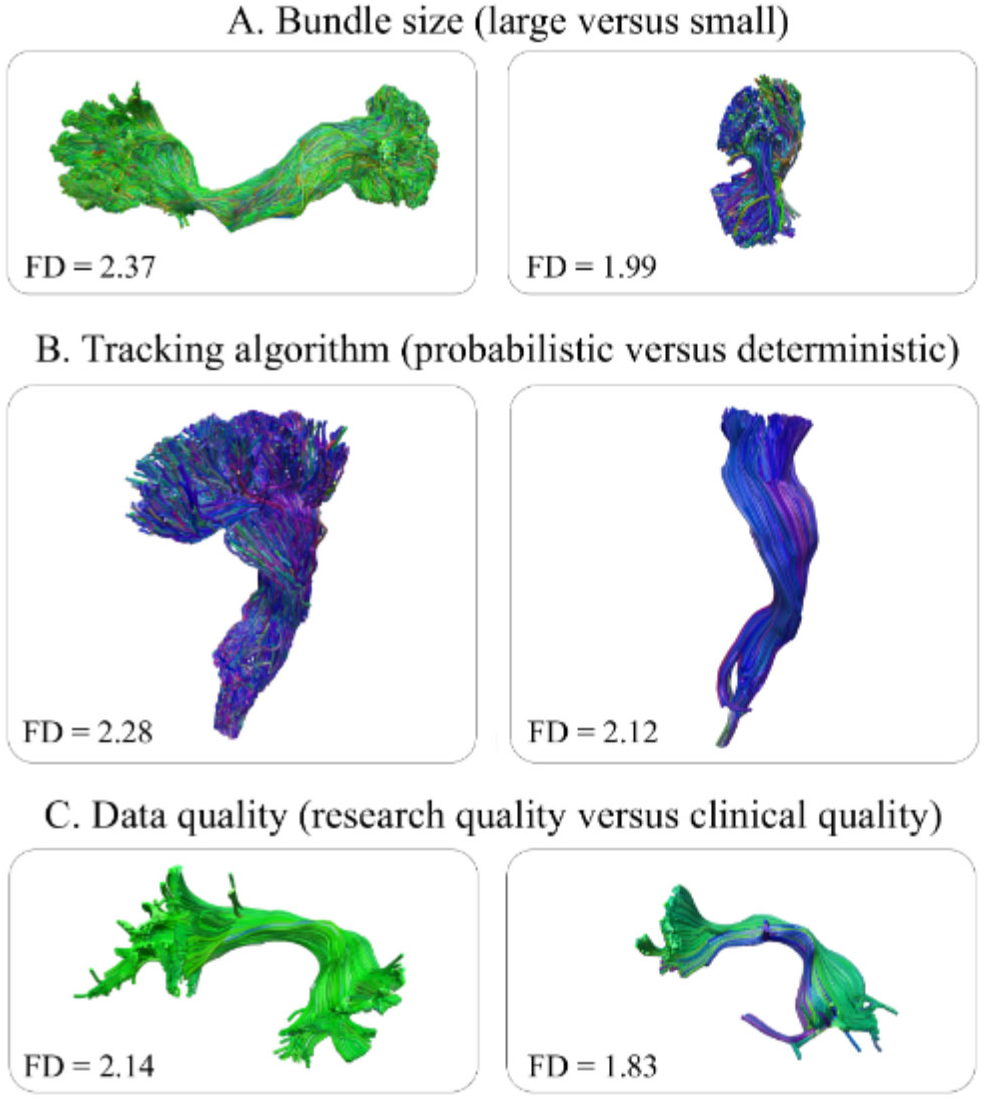
Examples of different properties of bundles. A. Two bundles with different size, on the left a large bundle (inferior-frontooccipital fascicle) and on the right a small bundle (posterior arcuate). B. Two bundles (corticospinal tracts) obtained using different tracking algorithms, on the left with probabilistic and on the right with deterministic tracking. C. Two bundles (arcuate fascicles) segmented from diffusion data of different quality, on the left at research quality and on the right at clinical quality. In each panel it is reported the fractal dimension (FD) of the voxel mask of the respective bundle.

As of today, no single method for bundle segmentation has been demonstrated to be robust, to bundle size, tracking method and data quality. The choice of the most appropriate pipeline for tractography is not unequivocal, but rather is strongly affected by the quality of the available diffusion Magnetic Resonance Imaging (dMRI) data, and changes according to the specific application, depending on the desired level of sensitivity/specificity (Thomas et al. (2014)). Similarly, even though the interest in large bundles is well established in multiple applications (Wandell (2016); Pestilli (2018)), small and short bundles, which we here call *minor* bundles, have recently received increasing attention, see Guevara et al. (2011); Wu et al. (2016b); Guevara et al. (2017); Bullock et al. (2019). For example, the relatively smaller bundles connecting the human dorsal and posterior cortices have been recently proven to be of great help in understanding how information flows in the human brain (Wu et al. (2016b); Bullock et al. (2019); Sani et al. (2019)). For these reasons, we believe that automatic methods for white matter bundle segmentation must be able to maintain a high quality of results across different settings.

The main contribution of the present work is a novel method for bundle segmentation that is robust to all properties described in Figure 1. We call the method *Classifyber*. Classifyber is a supervised streamline-based method, and is based on a linear classification model that predicts whether or not individual streamlines belong to the bundle of interest. It combines the current knowledge in bundle segmentation, exploiting both the similarity between streamlines, typical of streamline-based methods, and the anatomical information from ROIs, typical of connectivity-based methods. In contrast to state-of-the-art automatic segmentation methods, we claim that Classifyber is robust to different data settings.

As a second contribution, we present an extensive comparison between Classifyber and multiple other automatic bundle segmentation methods available in the literature, across a diverse set of conditions: major bundles vs minor bundles, different tractography techniques, and bundles from healthy subjects vs brain tumor patients. The results of these experiments support our claims that Classifyber is able to adapt to different data settings and sets a new standard with respect to the current literature by substantially improving the segmentation quality reached by other methods.

As a third contribution, we show that some segmentation methods are deeply affected by a geometrical property of the shape of the bundles: the *fractal dimension* (FD) (Zhang et al. (2006); Esteban et al. (2007)). Bundles with high fractal dimension are in general larger, more rounded, and have a smooth shape. Alternatively, bundles with low fractal dimension are generally smaller, flattened, and have a less smooth shape, see Figure 1. We observe that the tracking algorithm used to generate the tractography and the size of the bundles are among the main factors in the change of fractal dimension. The concept of the fractal dimension of a bundle is a key concept to discuss the experiments presented in this work.

This paper is structured as follows. In Section 2, we present the materials, which constitute four diverse as datasets, and describe the proposed method, Classifyber. In Section 3, we report the design and the results of a number of experiments that we conducted to verify our hypotheses. In Section 4, we discuss the results that suggest that practitioners should adopt the proposed Classifyber method as the leading standard for bundle segmentation.

## 2. Materials and Methods

In Section 2.1, we describe in detail the data materials used and those produced for this work. In Section 2.2 we present Classifyber, the proposed segmentation method.

### 2.1. Materials

In order to test different automatic bundle segmentation methods across a wide range of settings, we conducted extensive experiments across four different datasets of tractograms and bundles, three of which are novel. The description of these datasets, which we denote as HCP-minor, HCP-IFOF, HCP-major and Clinical, is provided in the following sections, together with the atlases used to derive the ROIs for the proposed method.

#### 2.1.1. Data sources

The first three datasets are built on top of diffusion data freely available from the Human Connectome Project (HCP) (Van Essen et al. (2013); Sotiropoulos et al. (2013)), 3T scanner, image resolution of 1.25 mm isotropic, 270 gradient directions with *b*-values=1000, 2000, and 3000 *s/mm*^2^ and 18 volumes with *b*=0. Data have already been preprocessed with the minimal pipeline of Glasser et al. (2013), which includes brain extraction and correction for motion, distortion and eddy-currents. The fourth dataset is an in-house clinical dataset built from patients with brain tumors, 1.5T scanner, image resolution 0.9 x 0.9 x 2.4 mm, 60 gradient directions with *b*-value=1000 *s/mm*^2^ and 1 volume with *b*=0. Data were corrected for eddy-current and motion, and an additional step of rescaling was applied to obtain an isotropic voxel resolution of 2 mm.

#### 2.1.2. Datasets of tractograms and expert-based segmented bundles

A comprehensive description of the four different datasets considered in this work is given in Table 1 and can be summarized as follows:

i. **HCP-minor**. Probabilistic ensemble tractograms with 750K streamlines of 105 HCP subjects with a collection of 8 minor bundles validated by expert neuroanatomists, such as the posterior arcuate fasciculus (pArc), see Bullock et al. (2019).
ii. **HCP-IFOF**. Deterministic tractograms with approximately 500K streamlines of 30 HCP subjects, with manual segmentations by one expert neurosurgeon of the inferior fronto-occipital fasciculus (IFOF).
iii. **HCP-major**. Probabilistic tractograms with 10 million streamlines of 105 HCP subjects, with a collection of major bundles, such as the corticospinal tract (CST) and the arcuate fasciculus (AF), segmented through a semi-automatic procedure, see Wasserthal et al. (2018a).
iv. **Clinical**. Deterministic tractograms with approximately 100K streamlines of 10 patients with brain tumors, with segmented IFOF and AF in the lesioned hemisphere, manually delineated by one expert neurosurgeon.

**Table 1:**
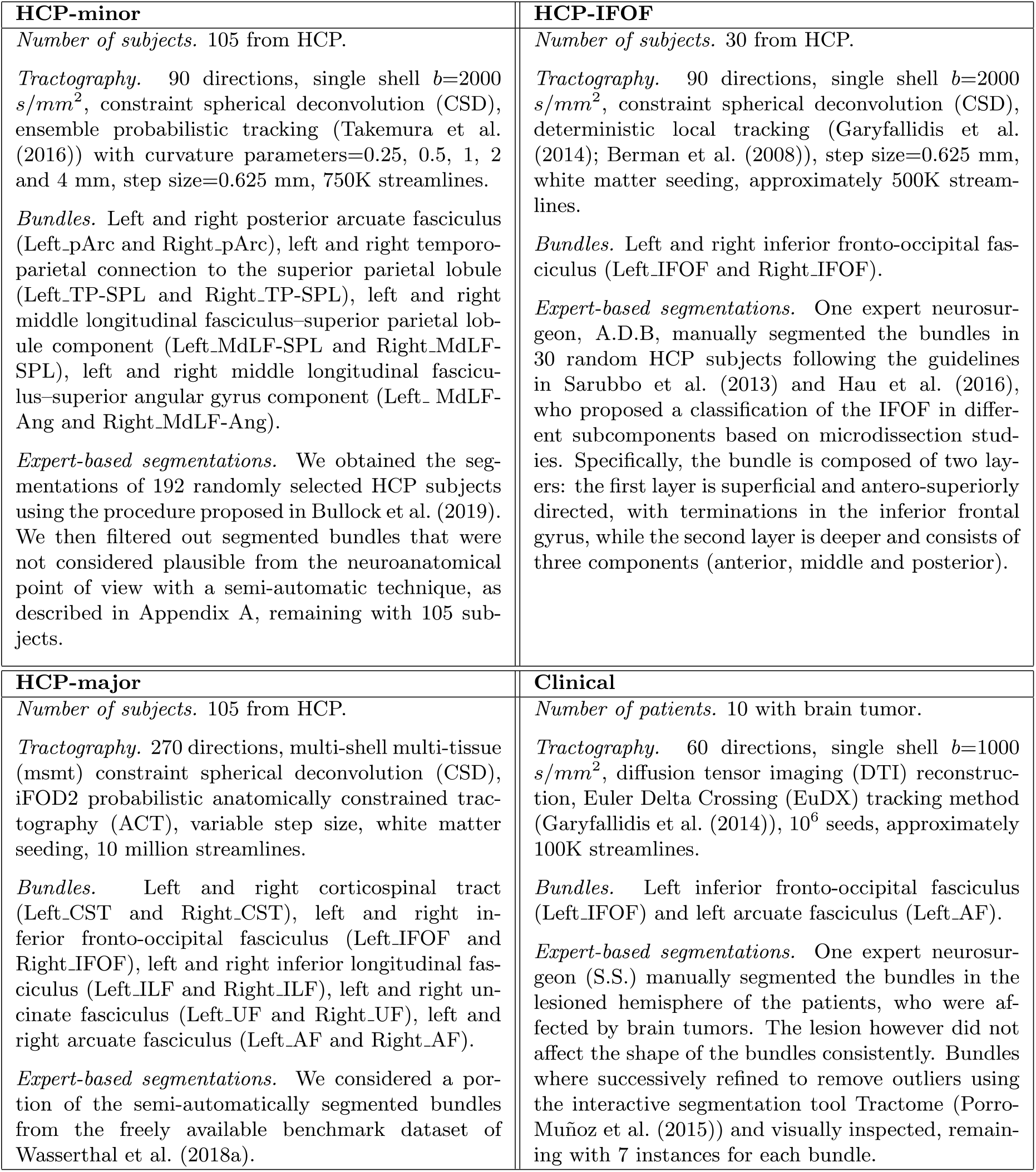
Comprehensive description of the four datasets of tractograms and expert-based segmented bundles, i.e. HCP-minor, HCP-IFOF, HCP-major and Clinical datasets.

|  |  |
| --- | --- |
| **HCP-minor**<br>*Number of subjects.* 105 from HCP.<br>*Tractography.* 90 directions, single shell $b=2000$ $s/mm^2$, constraint spherical deconvolution (CSD), ensemble probabilistic tracking (Takemura et al. (2016)) with curvature parameters=0.25, 0.5, 1, 2 and 4 mm, step size=0.625 mm, 750K streamlines.<br>*Bundles.* Left and right posterior arcuate fasciculus (Left_pArc and Right_pArc), left and right temporo-parietal connection to the superior parietal lobule (Left_TP-SPL and Right_TP-SPL), left and right middle longitudinal fasciculus–superior parietal lobule component (Left_MdLF-SPL and Right_MdLF-SPL), left and right middle longitudinal fasciculus–superior angular gyrus component (Left_MdLF-Ang and Right_MdLF-Ang).<br>*Expert-based segmentations.* We obtained the segmentations of 192 randomly selected HCP subjects using the procedure proposed in Bullock et al. (2019). We then filtered out segmented bundles that were not considered plausible from the neuroanatomical point of view with a semi-automatic technique, as described in Appendix A, remaining with 105 subjects. | **HCP-IFOF**<br>*Number of subjects.* 30 from HCP.<br>*Tractography.* 90 directions, single shell $b=2000$ $s/mm^2$, constraint spherical deconvolution (CSD), deterministic local tracking (Garyfallidis et al. (2014); Berman et al. (2008)), step size=0.625 mm, white matter seeding, approximately 500K streamlines.<br>*Bundles.* Left and right inferior fronto-occipital fasciculus (Left_IFOF and Right_IFOF).<br>*Expert-based segmentations.* One expert neurosurgeon, A.D.B, manually segmented the bundles in 30 random HCP subjects following the guidelines in Sarubbo et al. (2013) and Hau et al. (2016), who proposed a classification of the IFOF in different subcomponents based on microdissection studies. Specifically, the bundle is composed of two layers: the first layer is superficial and antero-superiorly directed, with terminations in the inferior frontal gyrus, while the second layer is deeper and consists of three components (anterior, middle and posterior). |
| **HCP-major**<br>*Number of subjects.* 105 from HCP.<br>*Tractography.* 270 directions, multi-shell multi-tissue (msmt) constraint spherical deconvolution (CSD), iFOD2 probabilistic anatomically constrained tractography (ACT), variable step size, white matter seeding, 10 million streamlines.<br>*Bundles.* Left and right corticospinal tract (Left_CST and Right_CST), left and right inferior fronto-occipital fasciculus (Left_IFOF and Right_IFOF), left and right inferior longitudinal fasciculus (Left_ILF and Right_ILF), left and right uncinate fasciculus (Left_UF and Right_UF), left and right arcuate fasciculus (Left_AF and Right_AF).<br>*Expert-based segmentations.* We considered a portion of the semi-automatically segmented bundles from the freely available benchmark dataset of Wasserthal et al. (2018a). | **Clinical**<br>*Number of patients.* 10 with brain tumor.<br>*Tractography.* 60 directions, single shell $b=1000$ $s/mm^2$, diffusion tensor imaging (DTI) reconstruction, Euler Delta Crossing (EuDX) tracking method (Garyfallidis et al. (2014)), $10^6$ seeds, approximately 100K streamlines.<br>*Bundles.* Left inferior fronto-occipital fasciculus (Left_IFOF) and left arcuate fasciculus (Left_AF).<br>*Expert-based segmentations.* One expert neurosurgeon (S.S.) manually segmented the bundles in the lesioned hemisphere of the patients, who were affected by brain tumors. The lesion however did not affect the shape of the bundles consistently. Bundles were successively refined to remove outliers using the interactive segmentation tool Tractome (Porro-Muñoz et al. (2015)) and visually inspected, remaining with 7 instances for each bundle. |

#### 2.1.3. Data preprocessing

For the three HCP datasets, we computed the nonlinear warp to register the structural T1-weighted images of every subject of each dataset to the MNI152 T1 template using the Advanced Normalization Tool (ANTs) (Avants et al. (2008)). For the clinical dataset, we computed a streamline linear registration (SLR) to the whole brain template of Yeh et al. (2018)^1^, because non-linear registration of clinical data is debated, as reported in Garyfallidis et al. (2015). In both cases, we applied the registrations to tractograms and bundles.

#### 2.1.4. Atlases

We exploited the following freely available atlases in order to derive the ROIs used by Classifyber, which were then registered to the MNI152 T1 template.

##### MNI152_ICBM2009c_reconstructed_atlas

This atlas^2^, is a curated FreeSurfer parcellation of the ICBM2009c nonlinear asymmetric template, see Gorgolewski (2016) and Fonov et al. (2011). These ROIs are used to define the terminal regions of minor bundles.

##### MNI_JHU_tracts_ROIs_atlas

This atlas^3^ is composed of two planar waypoint ROIs for each of 20 major bundles, which delineate the path of each bundle before it diverges towards the cortex. Each ROI was drawn on a group-average dataset in MNI space, see Wakana et al. (2007).

### 2.2. Methods

Classifyber is a novel method that performs automatic bundle segmentation as a supervised learning problem, meaning that the algorithm learns how to segment from expert-based examples. The name *Classifyber* is the linguistic blend of *Classify* and *fiber*, which explains the basic principle of its functioning: to classify whether or not a given streamline/fiber^4^ belongs to the bundle of interest.

Below, we provide a formal description of Classifyber, from the basic concepts to the key element of the proposed method, i.e., the vectorial representation of a streamline that merges geometrical information typically used by streamline-based segmentation methods, and anatomical information, typically used by connectivity-based segmentation methods. We conclude the section by introducing the notion of the *fractal dimension* (FD) of a bundle, which will be used to discuss the experimental results in Section 4.

#### 2.2.1. Basic concepts

A *streamline s* = [***x***_1_, …, ***x***_*n*_] is an ordered sequence of 3D points, **x**_*i*_ = [*x*_*i*_, *y*_*i*_, *z*_*i*_] ∈ ℝ^3^, *i* = 1 … *n*, that approximates a group of axons with a similar path in the white matter of the brain. A *tractogram T* is the entire set of streamlines of the white matter of a brain: *T* = {*s*_1_, …, *s*_*M*_}, where *M* typically ranges from hundreds of thousands to several millions. A white matter *bundle*, *b ⊂ T*, is a subset of the tractogram with a specific anatomical meaning, such as the corticospinal tract.

Experts neuroanatomists manually segment a given bundle *b* in a tractogram adopting several strategies, which may comprise the definition of inclusion/exclusion ROIs to obtain the desired streamlines. From the point of view of an algorithm, a convenient way to model that segmentation process is to consider each streamline individually and to decide whether or not the streamline belongs to the bundle:

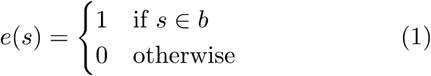

where *e*(*s*) denotes the expert deciding on the streamline *s*.

#### 2.2.2. Classifyber

Classifyber implements a classifier that accurately predicts whether or not a given streamline *s* belongs to the bundle *b*. In analogy to the previous work of Olivetti and Avesani (2011), in this work we propose a linear classifier method as core algorithm for Classifyber, for multiple reasons: it is extremely well known and easy to understand, it is very fast and requires minimal resources, software implementations are commonly available and, as opposed to non-linear methods, it can be interpreted. Generally, a linear classifier *l* takes as input a vector of real values **v** ∈ ℝ^*d*^ and returns its predicted category, i.e. 0 or 1:

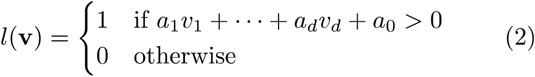

where the weights of the linear classifier *a*_0_, …, *a*_*d*_ are estimated by minimizing the errors in classification on a training set (plus regularization terms that may differ between different algorithms).

In order to use a linear classifier on a streamline *s*, it is necessary to transform the streamline into a vector *v* which contains the necessary information for the task of bundle segmentation. In other words, we need to define an effective *feature space* to represent streamlines as vectors. This is a key step in the proposed method, where we extract geometrical and anatomical information from the streamline and create this vector.

The proposed feature space is based on the general concept of *dissimilarity representation*, see Pekalska and Duin (2005) and Olivetti et al. (2012), which states that an object, i.e. a streamline, can be accurately represented by its distances from a fixed set of objects called *landmarks*, i.e. a fixed set of streamlines, that acts as reference system. The feature space is composed of four parts, which we describe here in an intuitive way and illustrate in Figure 2. Two parts, set 1 and set 3, are composed of global features independent from the definition of bundle of interest, while the other two parts, set 2 and set 4, are composed of bundle-specific features. We provide a comprehensive description of the procedure adopted to build the feature space in Appendix B.1. Given a streamline *s*, the four sets of feature values are:

i. **Set 1: streamline distances from 100 *global* landmarks.** The global landmarks are 100 streamlines evenly spread over a whole tractogram. The minimum average direct flip distance (*d*_MDF_) is one of the most commonly adopted distance function between streamlines, see Garyfallidis et al. (2012) and Olivetti et al. (2017).
ii. **Set 2: streamline distances from 100 *local landmarks*.** The local landmarks are 100 streamlines evenly spread locally to the area of the bundle.
iii. **Set 3: endpoint distance from 100 *global* landmarks.** This part of the feature space describes the anatomical connectivity pattern of the streamlines. The *endpoint* distance (*d*_END_) is the distance between endpoints of the two streamlines, because two streamline with neighboring endpoints connect the same areas, see Bertò et al. (2019).
iv. **Set 4: 2 ROI distances.** All bundles considered in this work can be normatively defined through two cortical ROIs or two waypoint ROIs. The ROI distance between a streamline and an ROI (*d*_ROI_) is the minimum among the distances between the points of the streamline and the voxels of the ROI, see Bertò et al. (2019).

**Figure 2:**
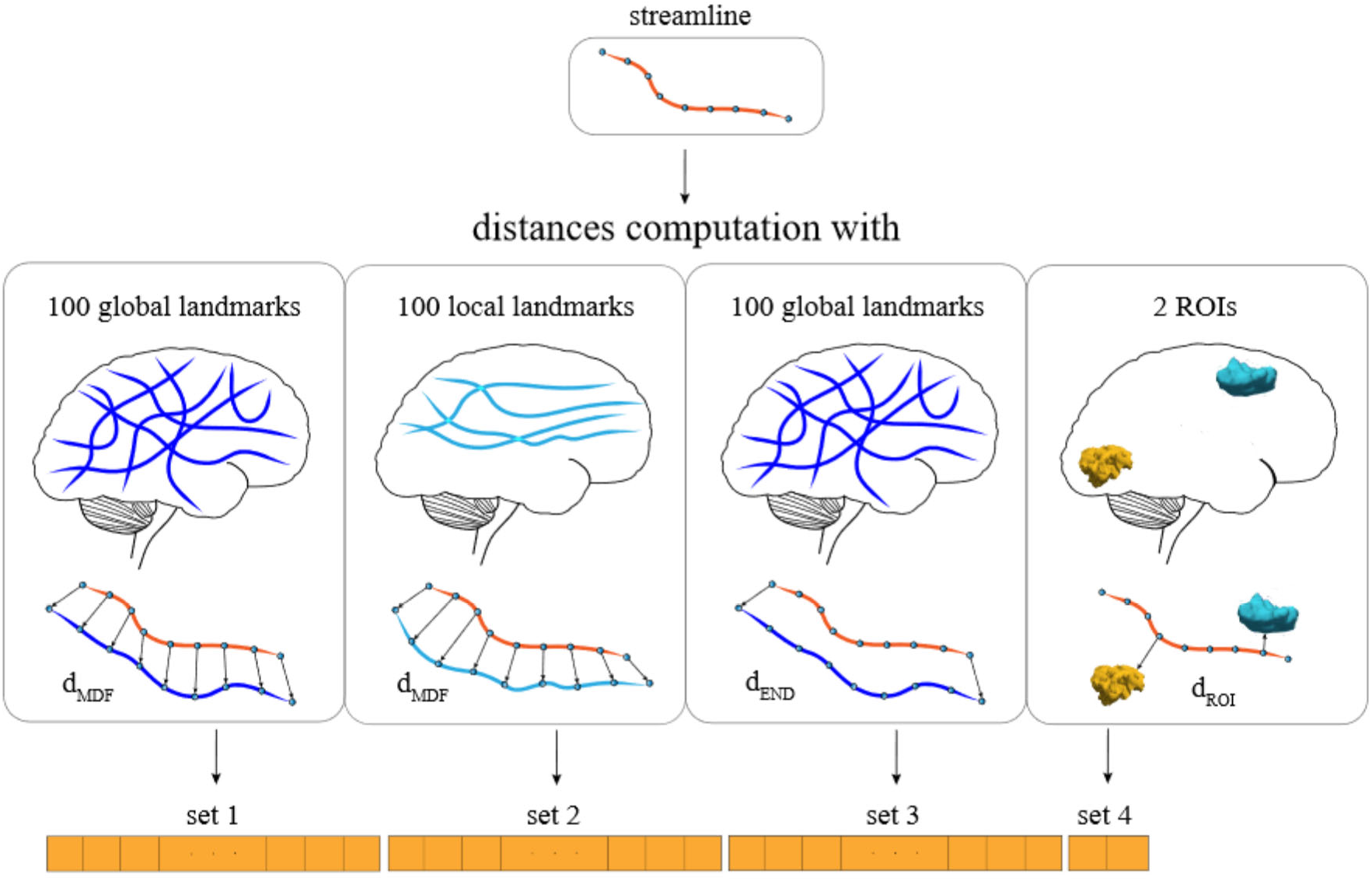
Feature definition and extraction. Set 1 and set 2 contain the distances (*d*_MDF_) of the streamline with 100 global and 100 local landmarks, respectively. Set 3 contains the distances (*d*_END_) between the endpoints of the streamline and the 100 global landmarks. Set 4 contains the distances (*d*_ROI_) between the streamline and the two ROIs pertaining to the bundle of interest.

Overall, the proposed feature space consists of *d* = 302 features, meaning that each streamline can be transformed into a vector of 302 values.

Therefore, given the tractograms and the expert-based segmented bundles of Section 2.1 of multiple subjects, we first transform all streamlines into vectors and label them with 1 or 0, to indicate whether or not they belong to the bundle of interest, and then train a classifier to segment a specific bundle, e.g. the corticospinal tract (CST). Notice that, in order to segment different kinds of bundles, it is necessary to train different instances of Classifyber, each with a set of examples of the desired kind of bundle. Afterwards, given a tractogram of a new subject, we predict whether or not the streamlines belong to that bundle. This procedure, which we divided into a *training phase* and a *test phase*, is summarized below and illustrated in Figure 3. A more comprehensive description is given in Appendix B.2.

**Figure 3:**
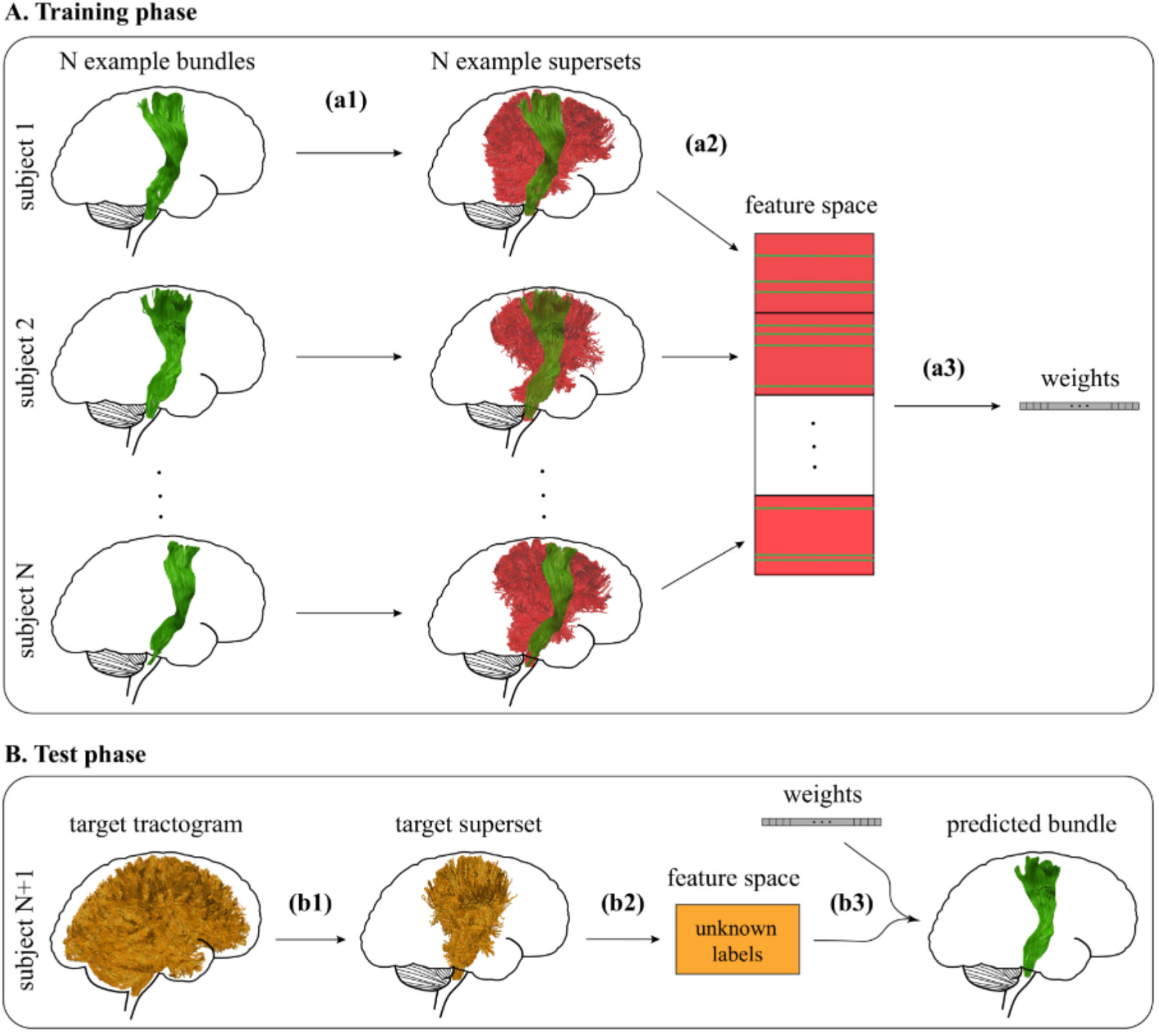
Training and test of Classifyber. A. Schematic illustration of the training phase of Classifyber for a given bundle (CST) over *N* different subjects. Step (a1): bundle superset. Streamlines belonging to the bundle are depicted in green (class 1), while those not belonging to the bundle are depicted in red (class 0). Step (a2): feature extraction. Step (a3): training of a linear Logistic Regression (LR) classifier. The outcome of this phase is a vector of weights. B. Schematic illustration of the test phase of Classifyber on a single target subject. Step (b1): bundle superset. All the streamlines are depicted in orange because the labels are unknown. Step (b2): feature extraction. Step (b3): test using the resulting weights of the training phase. The outcome of this phase is the predicted bundle (CST) in the target subject.

#### 2.2.3. Classifyber: training phase

The training phase is composed of three steps, which are schematically illustrated in Figure 3 (A).

##### Step (a1) *Bundle superset*

We reduce the number of streamlines in the training set by considering only the streamlines of the example bundles and their neighboring streamlines. This step helps to increase the classification accuracy and to reduce the computational cost of the next steps.

##### Step (a2) *Feature extraction*

Each streamline of the superset is then transformed into a vector, as described in Section 2.2.2. To the vector is assigned a class label 1 if it belongs to the bundle, 0 otherwise, see Figure 3 (A), where they are represented in green and red respectively. Step (a3) *Training*. A binary Logistic Regression classifier is trained using the fast stochastic average gradient (SAG) solver (Schmidt et al. (2017)), which returns the vector of weights *a*_0_, …, *a*_*d*_.

#### 2.2.4. Classifyber: test phase

The test phase is performed on one subject of the test set at the time, called the *target subject*. Similarly to the training phase, the test phase comprises three steps, which are schematically illustrated in Figure 3 (B).

##### Step (b1) *Bundle superset*

We reduce the whole target tractogram to a superset of the target bundle, which we estimate from training examples. The main reason for this step is to reduce the computational cost of the next steps.

##### Step (b2) *Feature extraction*

Each streamline of the bundle superset is then transformed into a vector, as described in Section 2.2.2. In this case, the class labels are unknown.

##### Step (b3) *Test*

By exploiting the linear classifier obtained from the training phase in step (a3), each streamline of the superset is predicted to be either part of the bundle or not, obtaining the predicted bundle in the target subject.

#### 2.2.5. Other bundle segmentation methods

In Section 3.1 we compare Classifyber to state-of-theart automatic segmentation methods. We selected two methods based on the recent extensive comparison presented in Wasserthal et al. (2018a), where *TractSeg* obtained the highest quality of bundle segmentation and *RecoBundles* ranked as the second best method among those freely available. In our comparison we also included *LAP*, see Sharmin et al. (2018), because it was not compared in Wasserthal et al. (2018a) but proved to be superior to nearest neighbor methods, the category to which RecoBundles belongs. In some cases, we used variants of TractSeg and RecoBundles, referred to as *TractSeg-retrained* and *RecoBundles-atlas*. We provide details on these other segmentation methods in Section 3.1.2.

#### 2.2.6. Evaluation procedure

To quantitatively evaluate the performance of the different segmentation methods we use a procedure commonly adopted in this literature, see for example Garyfallidis et al. (2018); Sharmin et al. (2016); Wasserthal et al. (2018a). We compute the degree of voxel overlap between the automatically segmented bundle 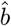 and the expert-based segmented bundle *b*, through the Dice Similarity Coefficient (DSC) (Dice (1945)): 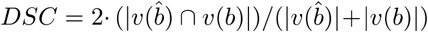 where |*v*()| is the number of voxels in the mask. The DSC ranges from 0 to 1 and the closer the score is to 1, the more the two bundles 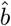 and *b* are similar. The evaluation is conducted in the subject’s native space.

#### 2.2.7. Fractal dimension

The concept of *fractal dimension* (FD) (Mandelbrot (1982)) can be used to quantify the degree of irregularity of a 3D shape. This notion has already been applied to the shape of the brain white matter (Zhang et al. (2006)) and to characterize multiple sclerosis (Esteban et al. (2007)).

Intuitively, for standard objects like straight lines, a 2D flat square or a 3D cube, the FD is 1, 2 and 3, respectively. Irregular lines can have FD greater than 1 and asymptotically 2, if their resulting shape is close to a 2D surface. In the same way, a convoluted 2D shape that resembles a 3D shape, or a 3D shape with several holes, both have FD between 2 and 3. For example, Zhang et al. (2006) estimated the FD of the 3D voxel mask of the white matter of human brains and obtained values between 2.1 and 2.5.

In this work, we determine the FD of the voxel mask of white matter bundles via the box-counting dimension, see Falconer (2014). The box-counting dimension is based on the idea of covering a given shape with boxes of size *σ* and it quantifies how the number of boxes changes when *σ* changes, in double-log scale:

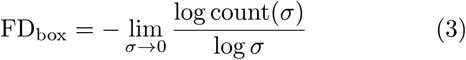

where count(*σ*) is the number of the necessary boxes. As an example, see the FD of some bundles in Figure 1.

#### 2.2.8. Data and code availability

We provide the source code of Classifyber and the code to estimate the box-counting dimension (with examples) as open source software, see Table 2. Moreover, Classifyber can be freely used as web application on the online platform `brainlife.io`. In Table 2 we list the web links related to all the implementations of the bundle segmentation methods considered in this work. Moreover, we freely share tractograms and expert-based segmented bundles of the HCP-minor dataset through the `brainlife.io` platform at https://doi.org/10.25663/brainlife.pub.11. The HCP-major dataset is available at https://doi.org/10.5281/zenodo.1477956. The HCP-IFOF is available upon formal data sharing agreement with the authors. The access to the Clinical dataset is limited by ethical and privacy issues and requires formal agreement with the neurosurgery unit involved in this study.

**Table 2:** Where to find the code and web apps of the methods considered in this work.

| code / web app | web link | contribution of this work |
| --- | --- | --- |
| Classifyber code | https://github.com/FBK-NILab/app-classifyber | yes (original) |
| Classifyber web app | https://doi.org/10.25663/brainlife.app.228<br>https://doi.org/10.25663/brainlife.app.265 | yes (original)<br>yes (original) |
| TractSeg code | https://github.com/MIC-DKFZ/TractSeg | no, already available |
| TractSeg web app | https://doi.org/10.25663/brainlife.app.186 | no, already available |
| TractSeg-retrained web app | https://doi.org/10.25663/brainlife.app.204<br>https://doi.org/10.25663/brainlife.app.205 | yes (adapted)<br>yes (adapted) |
| RecoBundles(-atlas) code | http://nipy.org/dipy | no, already available |
| LAP web app | https://doi.org/10.25663/brainlife.app.209 | yes (adapted) |
| Box-counting dimension | https://github.com/FBK-NILab/fractal_dimension | yes (original) |

## 3. Experiments and Results

The results described below compare Classifyber with other segmentation methods to demonstrate that the proposed one substantially increases the quality of segmentation with respect to other methods, consistently across different data settings. To this end, in Section 3.1, we describe the experimental design of Classifyber and of the other state-of-the-art methods considered in this work, followed by the experiments conducted on the fractal dimension of bundles. To conclude, in Section 3.2, we report the results of all the experiments.

### 3.1. Experiments

The experiments were conducted on the four datasets described in Section 2.1: HCP-minor, HCP-major, HCPIFOF and Clinical. For each dataset, the entire pool of subjects was randomly divided into two groups: the *training set* and the *test set*. Bundles of the training set were used as examples to learn from, while bundles of the test set were used to assess the performance of the different methods. Notice that the exact same test sets were kept for all the methods compared. In this way, we could compare both the quality of segmentation obtained by each method averaged over the pool of test subjects, such as in an *unpaired* test, and the subject-by-subject comparison in segmenting each bundle, such as in a *paired* test, e.g. how frequently one method obtained better quality of segmentation than another method.

#### 3.1.1. Classifyber: experimental setup

We retrieved the ROIs pertaining to each bundles in order to build the feature space of Classifyber, using the available atlases described in Section 2.1.4. For the dataset HCP-minor, the two ROIs considered for each bundle are the two terminal ROIs, i.e. the cortical regions that the bundle of interest connects, derived from Bullock et al. (2019). Specifically, the MdLF-Ang and MdLF-SPL connect the parietal region to the lateral-temporal region, while the TP-SPL and pArc connect the parietal region to the temporal region. Each region was built by merging specific cortical parcellations of the *MNI152_ICBM2009c_reconstructed_atlas*. For the other three datasets, the ROIs considered are the two planar waypoint ROIs defined in the *MNI_JHU_tracts_ROIs_atlas*, see Wakana et al. (2007).

##### HCP-minor

We considered only subjects for which all bundles received an expert-made score of 3 or higher, according to the procedure explained in Section Appendix A, resulting in a set of 40 subjects. We randomly split this pool of subjects into a group of 15 for training and a group of 25 for testing. Additionally, within this dataset, we also studied how much the quality of segmentation of Classifyber was affected when changing the number of subjects in the training set from 1 to 60. In this case, we considered also subjects for which all bundles received an expert-made score of at least of 2.

##### HCP-IFOF

We randomly split the pool of subjects into a group of 15 for training and a group of 15 for testing.

##### HCP-major

For this dataset, which is part of the dataset used in Wasserthal et al. (2018a), we selected the same 21 test subjects used in the experiments presented there. In this way, we could directly compare our new results on the major bundles with theirs and, at the same time, we could test the reproducibility of their results. Of the 84 remaining subjects, 15 were randomly selected and used as training set for Classifyber. The kinds of bundles considered are those for which the two waypoint ROIs are available in the *MNI_JHU_tracts_ROIs_atlas*. In preliminary experiments, we observed that the 10 million streamlines of each tractogram in HCP-major were extremely redundant for training Classifyber and just using 10% of them, randomly selected, did not significantly change the results. By using just 10% of the streamlines we reduced the training time by a factor of 10 and the RAM usage by a factor of 4.

##### Clinical

Due to the small number of subjects in the dataset, instead of splitting the pool of 7 subjects into training and test sets, we ran Classifyber in two different ways: (i) we trained Classifyber on the IFOFs and AFs of the HCP-major dataset and then segmented the 7 patients in the Clinical dataset. We chose this dataset because it is part of the exact same dataset used for training Tract-Seg, to have fair comparison between the two methods. We refer to this case as *Classifyber-major*. (ii) We performed a cross-validation study with the leave-one-subject-out (LOSO) strategy, using *only* 6 subjects from the Clinical dataset as training set and the remaining subjects as test set, repeatedly. We refer to this case as *Classifyber-LOSO*. In this latter case we also aimed to show the ability of Classifyber to accurately segment bundles even when trained on a very small number of segmentations, in this case only 6. To conclude, for the IFOF, we ran one additional experiment where Classifyber was trained on the HCP-IFOF dataset. We refer to this case as *Classifyber-IFOF*.

#### 3.1.2. State-of-the-art methods: experimental setup

Here we describe the state-of-the-art automatic segmentation methods that we considered in our comparison, their necessary variants to experiment on all datasets, and their experimental setup.

##### TractSeg

TractSeg, a voxel-based method recently proposed by Wasserthal et al. (2018a), is based on fully convolutional neural networks (FCNNs) and segments 72 bundles simultaneously. Its output are the voxel masks of the segmented bundles. We adopted the openly available pretrained network, which was trained on 84 subjects, and tested it on the dMRI data of the target subjects. We used the default parameters and the postprocessing option, which removes holes and isolated voxels in the predicted voxel mask of the bundles.

##### TractSeg-retrained

When the bundle to be segmented was not available among those covered by TractSeg, we retrained the FCNN on new examples with a procedure discussed in a private communication with the authors of Wasserthal et al. (2018a). We refer to this variant as *TractSeg-retrained*. First, we trained a single FCNNs per dataset with default parameters, 250 epochs, fraction of validation subjects = 0.2 and data augmentation. Then, we tested the method enabling the postprocessing option. For the HCP-minor dataset, we trained the model both with the same 15 subjects used in the other methods, and also with 69 additional subjects by considering as well those subjects for which all bundles received a score of at least 2 (84 subjects in total). We provide evidence of the successful training in Appendix C.1.

##### RecoBundles-atlas

Garyfallidis et al. (2018) proposed a streamline-based segmentation method, called RecoBundles, that takes as input one example bundle which is used to estimate the corresponding bundle in a new tractogram by means of linear registration and nearest-neighboring streamlines. We contacted the authors of RecoBundles and received the indication to use the bundle models provided by the atlas of Yeh et al. (2018)^5^ as the example bundles and specifically 30 (out of 80) of them. We denote as *RecoBundles-atlas* this use of the RecoBundles algorithm. We used the best configuration of parameter values found from an extensive preliminary assessment analogous to the one reported in Appendix C.2. This configuration uses default parameter values with the exception of disabling the local streamline linear registration (SLR) option (most probably because all the datasets were already coregistered in MNI space) and using the minimum average mean distance (*d*_*MAM*_) instead of the minimum average direct flip distance (*d*_*MDF*_).

##### RecoBundles

When the bundle to be segmented is not available among the 30 selected bundles from the atlas of Yeh et al. (2018), we fell back to the original indication in Garyfallidis et al. (2018) and used the same example bundles adopted as input for the other methods. We denote this use of the algorithm plainly as *RecoBundles*. Due to the fact that RecoBundles accepts only one bundle as example, to quantify the quality of segmentation when multiple bundles are available in the training set, we adopted a procedure similar to the one used in the experiments of Wasserthal et al. (2018a). Specifically, we treated the *N* example bundles as models for *N* separate runs of the algorithm over the target subject, thus obtaining *N* different predictions of the same bundle. We then evaluated the segmentation accuracy by computing the mean DSC across the *N* bundles. As for RecoBundles-atlas, we used the best configuration of parameter values found from an extensive preliminary assessment described in Appendix C.2.

##### LAP

Sharmin et al. (2018) proposed a streamline-based segmentation method that takes as input multiple example bundles which are used to estimate the corresponding bundle in a target tractogram by means of finding corresponding streamlines through the solution of a Linear Assignment Problem (LAP) and a refinement step. We ran the algorithm following the original procedure and we set the parameter *k*, the only parameter of the method, corresponding to the number of nearest neighbors streamlines to compute the superset, equal to 2000 (default *k* = 500), since the total number of streamlines of the tractograms considered in our experiments are approximately 4 times higher than in the original study of Sharmin et al. (2018). One limitation of LAP is that it is computationally too expensive for supersets larger than 100 thousands streamlines, both for memory and time requirements.

#### 3.1.3. Experiments on Fractal Dimension (FD)

In this experiment, we studied how the performance of the different segmentation methods is affected by the FD of the target bundles. We computed the FD of the voxel mask of each target bundle as segmented by experts and compared it with the quality of segmentation (DSC) obtained for that bundle by each automatic segmentation method, across all experiments (approximately 500 bundles). For TractSeg and RecoBundles, that number was larger because we investigated also the variants TractSeg-retrained and RecoBundles-atlas, while for LAP it was smaller because it was not possible to execute the method on the HCP-major dataset, where supersets substantially exceeded 100 thousands streamlines.

### 3.2. Results

#### 3.2.1. Results on HCP-minor dataset

In Table 3 and Figure 6, we quantify the mean quality of segmentation in terms of DSC across the minor bundles considered in this set of experiments for RecoBundles, TractSeg-retrained, LAP and Classifyber across 25 subjects. TractSeg and RecoBundles-atlas were excluded because they do not address minor bundles. The quality of segmentation obtained by Classifyber is very high and outperforms all the other methods. Moreover, given that the target subjects are exactly the same across all methods, we can also summarize the results with a direct comparison on the individual bundles: over the 200 segmentations (8 different bundles for each of the 25 test subjects) performed by each method during the test phase, Classifyber obtained higher quality of segmentation (higher DSC) than RecoBundles and TractSeg-retrained in 100% of the cases, and than LAP in 99% of the cases.

**Table 3:** Quantitative comparison over HCP-minor dataset: DSC (mean sd) across 25 target subjects for RecoBundles, TractSeg-retrained, LAP and Classifyber. Highest quality of segmentation in bold face.

|  | RecoBundles | TractSeg-ret. | LAP | Classifyber |
| --- | --- | --- | --- | --- |
| Right_pArc | $0.76 \pm 0.04$ | $0.77 \pm 0.03$ | $0.80 \pm 0.03$ | **$0.88 \pm 0.03$** |
| Left_MdLF-Ang | $0.71 \pm 0.04$ | $0.72 \pm 0.06$ | $0.79 \pm 0.05$ | **$0.87 \pm 0.03$** |
| Left_pArc | $0.73 \pm 0.05$ | $0.75 \pm 0.03$ | $0.79 \pm 0.04$ | **$0.85 \pm 0.05$** |
| Right_MdLF-Ang | $0.68 \pm 0.04$ | $0.70 \pm 0.03$ | $0.76 \pm 0.03$ | **$0.84 \pm 0.03$** |
| Left_MdLF-SPL | $0.63 \pm 0.06$ | $0.67 \pm 0.05$ | $0.73 \pm 0.04$ | **$0.82 \pm 0.04$** |
| Right_TP-SPL | $0.62 \pm 0.08$ | $0.68 \pm 0.06$ | $0.72 \pm 0.05$ | **$0.82 \pm 0.05$** |
| Left_TP-SPL | $0.63 \pm 0.06$ | $0.67 \pm 0.04$ | $0.70 \pm 0.05$ | **$0.81 \pm 0.04$** |
| Right_MdLF-SPL | $0.60 \pm 0.05$ | $0.64 \pm 0.04$ | $0.70 \pm 0.03$ | **$0.80 \pm 0.04$** |

TractSeg-retrained, when trained on 84 subjects, performed better than TractSeg-retrained on 15 subject, obtaining a marginal increase in DSC between 0 and 0.03. For a fair comparison with the other methods, this result is not reported in Table 3 and Figure 6.

The superiority of Classifyber over the other segmentation methods is also evident from the qualitative comparison in Figure 4, in which the segmentations provided by the proposed method are, for all the bundles considered, the most anatomically similar to the expert-based segmentations. When using other methods, we observe a consistent bias in the predictions: RecoBundles and LAP tend to overestimate the bundle producing several false positives streamlines. On the other hand, for the majority of the bundles of this dataset, TractSeg-retrained correctly identifies the core part of the bundles, but fails to retrieve part of the cortical terminations. Illustrative examples of this behavior are in the last row of Figure 4, in which the Right MdLF-SPL is overestimated by RecoBundles (first panel), and it is missing most of the terminations in the latero-temporal ROI by TractSeg-retrained (second panel).

**Figure 4:**
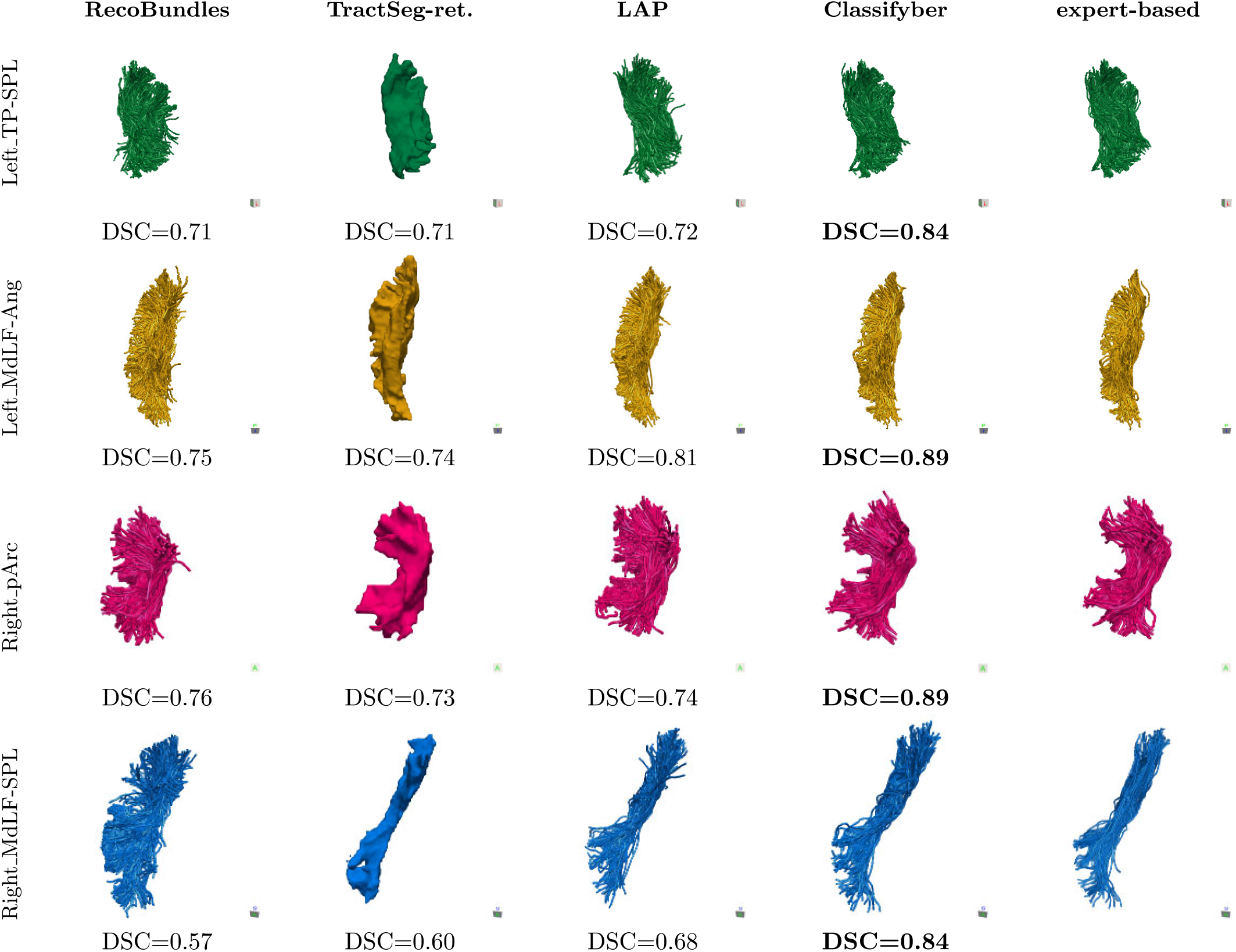
Qualitative comparison of segmented bundles in one target subject. Bundles on the rows: Left TP-SPL (first row), Left MdLF-Ang (second row) Right pArc (third row), and Right MdLF-SPL (fourth row). Automatic segmentation methods on the columns: RecoBundles (first column), TractSeg-retrained (second column), LAP (third column) and Classifyber (fourth column) and expert-based segmentation (fifth column). Highest quality of segmentation in bold face.

#### 3.2.2. Results on HCP-IFOF dataset

In Table 4 and Figure 6 we report the result of comparing Classifyber with all other methods and variants: RecoBundles-atlas, RecoBundles, TractSeg, TractSeg-retrained and LAP. The average DSC across 15 subjects shows the superiority of Classifyber. Moreover, in all individual cases, i.e. the 30 segmented bundles of the test set, Classifyber always obtained the highest DSC as compared to all other methods.

**Table 4:** Quantitative comparison over the HCP-IFOF dataset: DSC (mean sd) across 15 target subjects for RecoBundles-atlas, RecoBundles, TractSeg, TractSeg-retrained, LAP and Classifyber. Highest quality of segmentation in bold face.

|  | RecoBundles-atlas | RecoBundles | TractSeg | TractSeg-retrained | LAP | Classifyber |
| --- | --- | --- | --- | --- | --- | --- |
| Left_IFOF | $0.45 \pm 0.14$ | $0.80 \pm 0.04$ | $0.48 \pm 0.04$ | $0.61 \pm 0.03$ | $0.81 \pm 0.04$ | **$0.91 \pm 0.03$** |
| Right_IFOF | $0.62 \pm 0.18$ | $0.72 \pm 0.06$ | $0.41 \pm 0.06$ | $0.57 \pm 0.04$ | $0.73 \pm 0.05$ | **$0.89 \pm 0.03$** |

Additionally, a qualitative visual comparison is reported in Figure 5, which illustrates that the Left IFOF estimated with RecoBunldes-atlas (first panel), is clearly missing the middle and posterior subcomponents with respect to the expert-based segmented bundle (last panel). A very similar behavior is observed in the bundle predicted by TractSeg (third panel).

**Figure 5:**
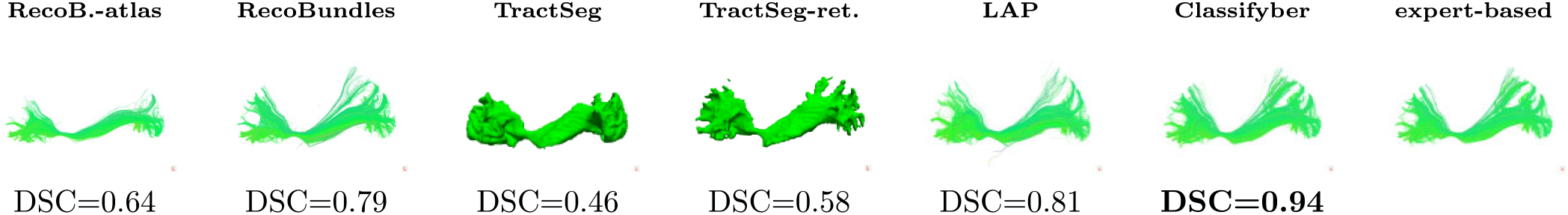
Qualitative comparison of segmented bundles in one target subject. One istance of Left IFOF for RecoBundles-atlas, RecoBundles, TractSeg, TractSeg-retrained, LAP and Classifyber with the expert-based segmented bundle. Highest quality of segmentation in bold face.

#### 3.2.3. Results on HCP-major dataset

In Table 5 and Figure 6 we report the mean quality of segmentation as DSC for RecoBundles-atlas, TractSeg and Classifyber over the major bundles considered, across 21 subjects. Over the 210 individual segmentations generated by each method in the test phase, Classifyber obtained a higher DSC than RecoBundles-atlas in 99% of the cases and higher than TractSeg in 86% of the cases.

**Figure 6:**
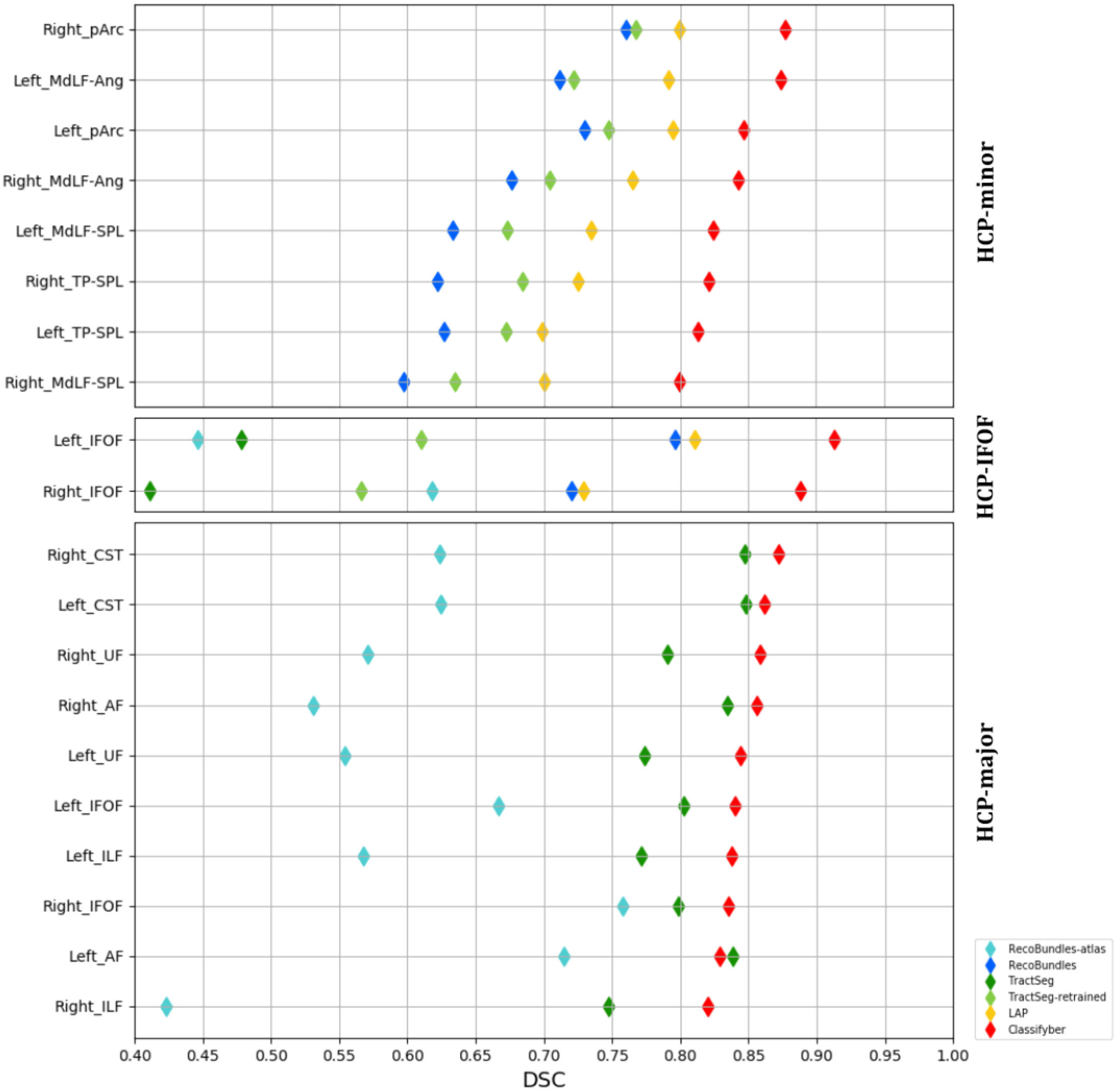
Summary of the quantitative comparison across the three HCP datasets. Top: mean DSC across 25 subjects of the HCP-minor dataset. Middle: mean DSC across 15 subjects of the HCP-IFOF dataset. Bottom: mean DSC across 21 subjects of the HCP-major dataset. The methods compared are depicted in different colors: RecoBundle-atlas (light blues), RecoBundles (blue), TractSeg (green), TractSeg-retrained (light green), LAP (yellow) and Classifyber (red).

**Table 5:** Quantitative comparison over HCP-major dataset: DSC (mean sd) across 21 target subjects for RecoBundles-atlas, TractSeg and Classifyber. Highest quality of segmentation in bold face.

|  | RecoB.-atlas | TractSeg | Classifyber |
| --- | --- | --- | --- |
| Right_CST | $0.62 \pm 0.07$ | **$0.85 \pm 0.02$** | **$0.87 \pm 0.02$** |
| Left_CST | $0.62 \pm 0.11$ | **$0.85 \pm 0.03$** | **$0.86 \pm 0.10$** |
| Right_UF | $0.57 \pm 0.24$ | $0.79 \pm 0.03$ | **$0.86 \pm 0.03$** |
| Right_AF | $0.53 \pm 0.11$ | **$0.83 \pm 0.02$** | **$0.86 \pm 0.03$** |
| Left_UF | $0.55 \pm 0.27$ | $0.77 \pm 0.03$ | **$0.84 \pm 0.04$** |
| Left_IFOF | $0.67 \pm 0.06$ | $0.80 \pm 0.02$ | **$0.84 \pm 0.03$** |
| Left_ILF | $0.57 \pm 0.07$ | $0.77 \pm 0.02$ | **$0.84 \pm 0.04$** |
| Right_IFOF | $0.76 \pm 0.04$ | $0.80 \pm 0.02$ | **$0.84 \pm 0.03$** |
| Left_AF | $0.71 \pm 0.05$ | **$0.84 \pm 0.03$** | **$0.83 \pm 0.04$** |
| Right_ILF | $0.42 \pm 0.13$ | $0.75 \pm 0.03$ | **$0.82 \pm 0.04$** |

#### 3.2.4. Results on Clinical dataset

In Table 6 we report the quantitative comparison in terms of mean DSC for Classifyber and TractSeg. The comparison is focused on TractSeg because in Wasserthal et al. (2018a) it is stated that the method is effective on clinical quality data as well, without the need for retraining the network. Individually, over the 14 segmented bundles, Classifyber always obtained a higher DSC than TractSeg, for all the different training sets, i.e. for all three different variants: Classifyber-major, Classifyber-IFOF and Classifyber-LOSO. In Figure 7 we show a qualitative comparison between the different cases.

**Table 6:** Quantitative comparison over the Clinical dataset: DSC (mean sd) across 7 target subjects for TractSeg, Classifyber-major, Classifyber-IFOF and Classifyber-LOSO. Highest quality of segmentation in bold face.

|  | TractSeg | Classifyber-major | Classifyber-IFOF | Classifyber-LOSO |
| --- | --- | --- | --- | --- |
| Left_IFOF | 0.42 ± 0.05 | 0.71 ± 0.09 | 0.82 ± 0.07 | **0.89 ± 0.03** |
| Left_AF | 0.23 ± 0.02 | 0.74 ± 0.13 | - | **0.92 ± 0.03** |

**Figure 7:**
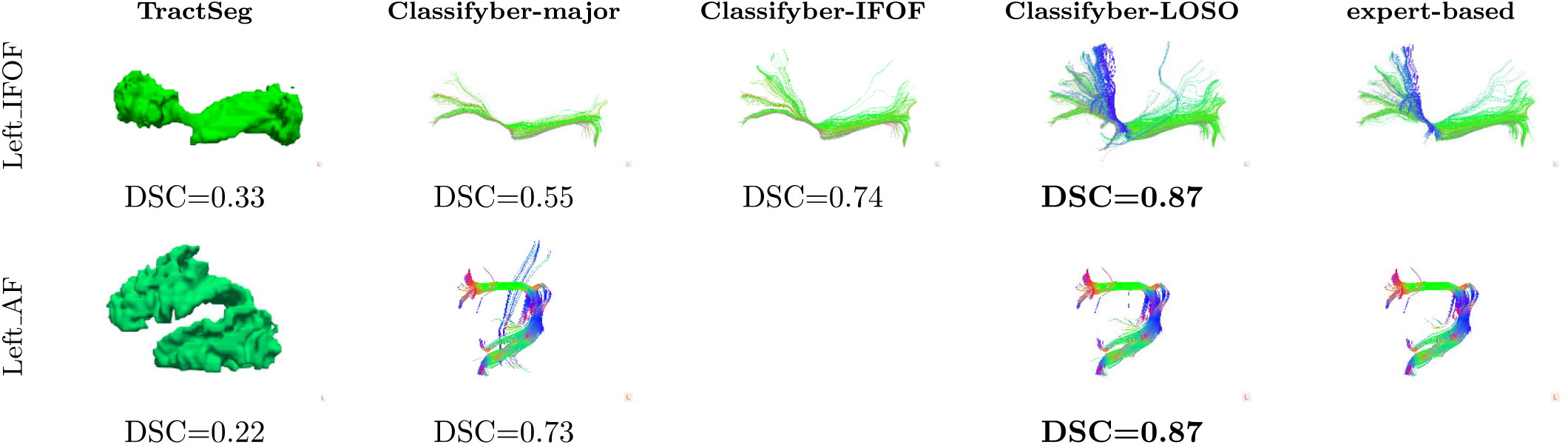
Qualitative comparison of segmented bundles in one of the patients. Bundles on the rows: Left IFOF (first row) and Left AF (second row). Automatic segmentation methods on the columns: TractSeg (first column), Classifyber-major (second column), ClassifyberIFOF (third column), Classifyber-LOSO (forth column) and expert-based segmentation (fifth column). Highest quality of segmentation in bold face.

#### 3.2.5. Results on Fractal Dimension (FD)

In Figure 8, we present the relationship between the FDs and the DSC scores of each method when segmenting all bundles in the experiments over the four datasets described above, i.e. on approximately 500 bundles. In the same figure, we also show the linear interpolation of such values as a summary of all experiments presented in this work, reporting the Pearson correlation coefficient (R) between FD and DSC. The results show that the quality of segmentation of TractSeg is strongly dependent on the FD of the bundle to be segmented. LAP also shows some degree of dependency, while RecoBundles and Classifyber are not affected by the FD of bundles. Additionally, in Table 7, we report the different range of FD values across the four datasets described in Section 2.1. Bundles of the HCP-major dataset have on average the highest FD, while bundles of the clinical dataset the lowest.

**Table 7:** FD values of the 4 datasets used in this work. The dataset are sorted according to their mean FD.

| dataset | FD min | FD max | FD mean ± sd |
| --- | --- | --- | --- |
| HCP-major | 2.09 | 2.44 | 2.30 ± 0.08 |
| HCP-minor | 1.89 | 2.26 | 2.10 ± 0.08 |
| HCP-IFOF | 1.74 | 2.08 | 1.99 ± 0.06 |
| Clinical | 1.75 | 1.96 | 1.86 ± 0.06 |

**Figure 8:**
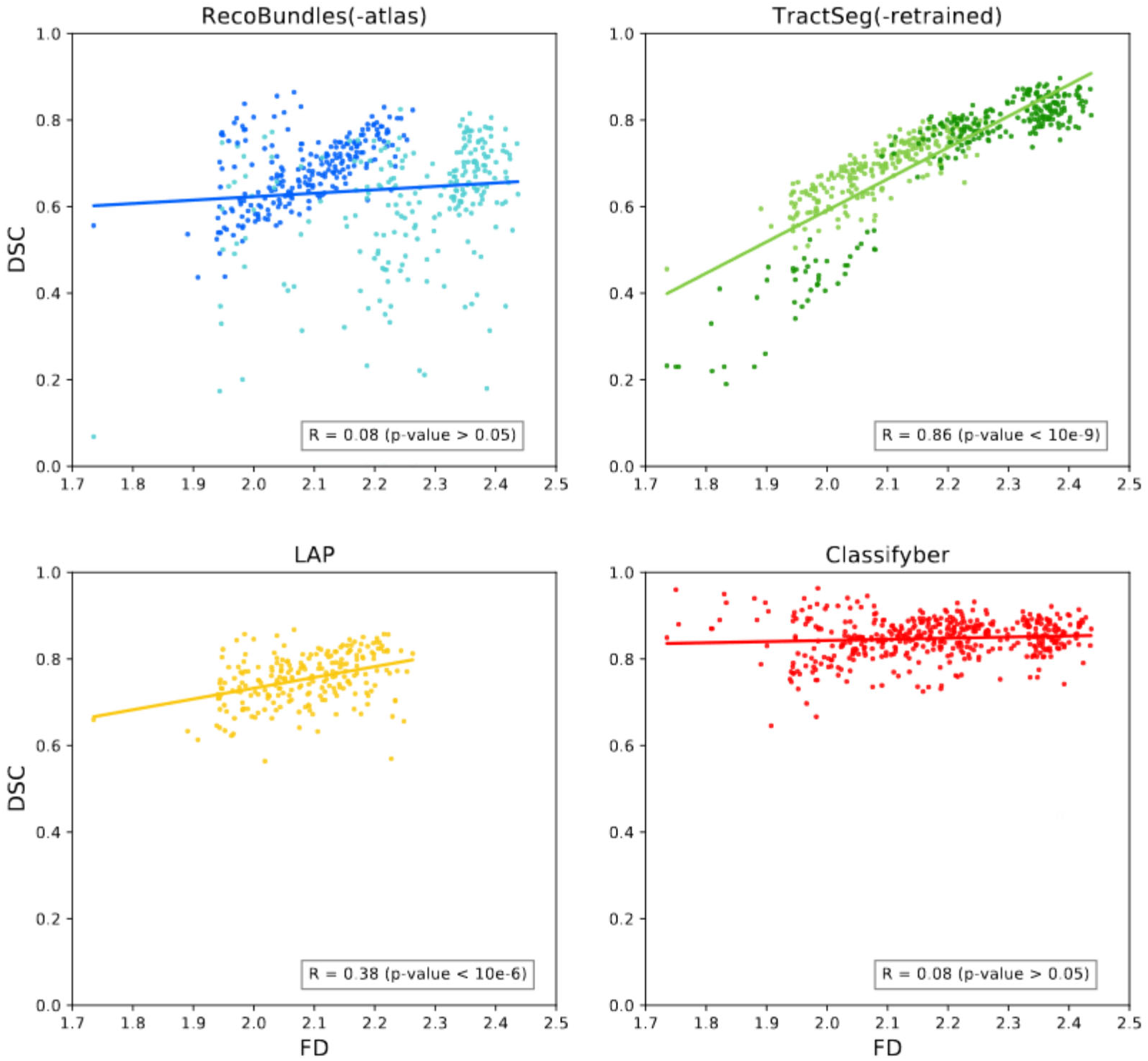
DSC vs FD across all methods for all the predicted bundles of the experiments in this work. From the left: RecoBundles and RecoBundles-atlas (blue and light blue), TractSeg and TractSeg-retrained (green and light green), LAP (yellow), and Classifyber (red). R is the Pearson correlation coefficient and related *p*-value between FD and DSC over all the predicted bundles, i.e. approximately 500 segmentations.

#### 3.2.6. Classifyber: the size of the training set

For all automatic segmentation methods that learn from examples, the higher the number of training subjects, the better the resulting quality of segmentation. Nevertheless, in practice, the cost of time and effort by an expert to prepare a curated training set severely limits this number. In Figure 9 we show how the mean DSC of Classifyber over multiple bundles changes with the number of training subjects. We observe that the quality of segmentation has no substantial increase beyond approximately 15 subjects and plateaus at 30 subjects.

**Figure 9:**
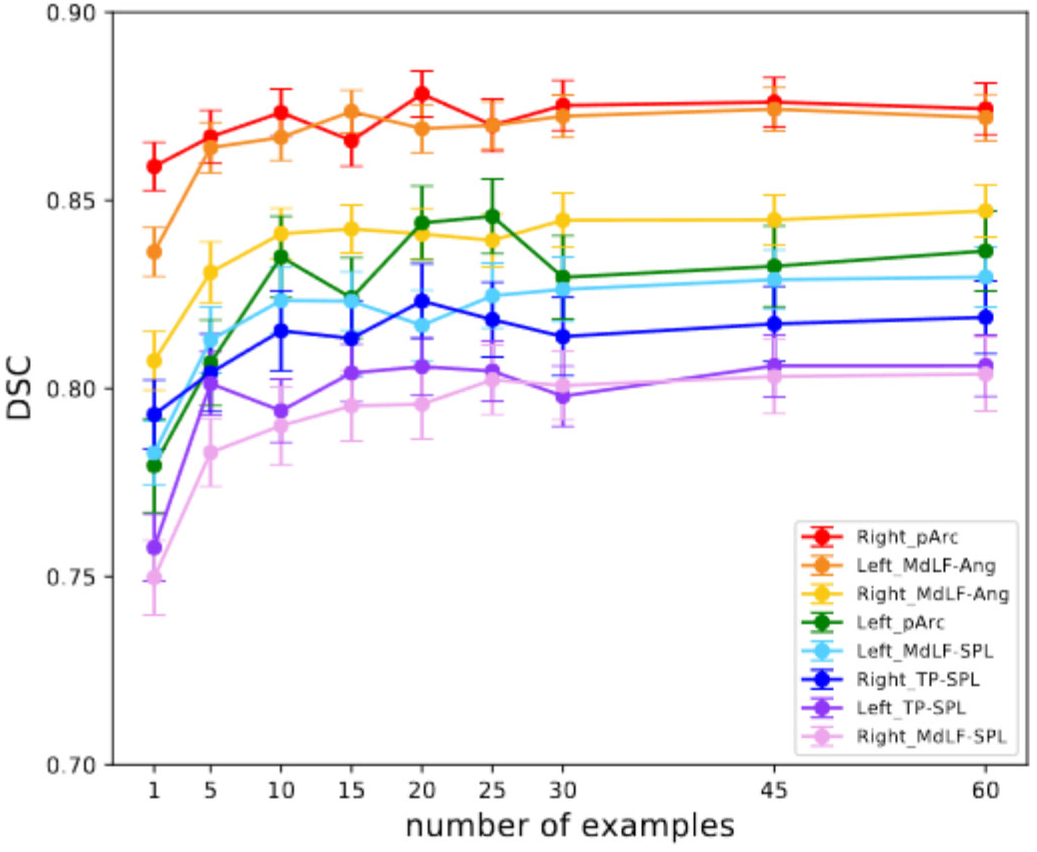
DSC (mean sd of the mean) across 25 test subjects of the HCP-minor dataset when varying the number of examples, from 1 to 60. Each of the bundle is depicted with a different color.

#### 3.2.7. Analysis of the computing time

In Table 8 we report the time required by each segmentation method for the training phase and for segmenting one bundle of the HCP-IFOF dataset. We chose this dataset because it is the only dataset on which we compared all segmentation methods and variants.

**Table 8:** Time in minutes required to train each method and to segment one IFOF for: RecoBundles-atlas, RecoBundles, TractSeg, TractSeg-retrained, LAP and Classifyber, when having 15 training examples. (*) GPU accelerated. (**) segmenting 2 kinds of bundles at the same time. (***) Training on 84 subjects to segment 72 kinds of bundles at the same time.

|  | training phase | segmentation | total |
| --- | --- | --- | --- |
| RecoBundles-atlas | 0 | 0.5 | 0.5 |
| RecoBundles | 0 | 3 | 3 |
| Classifyber | 34 | 3 | 37 |
| LAP | 0 | 130 | 130 |
| TractSeg-ret.(* ) | 175(**) | 5 | 180 |
| TractSeg(*) | 720(***) | 5 | 725 |

In Figure 10 we show how the 37 minutes indicated in Table 8, needed to train Classifyber and to segment one bundle, are partitioned across the 6 steps (3 for the training phase and 3 for the test phase) described in Section 2.2.3 and Section 2.2.4. We observed that the training time is linearly correlated with the number of training streamlines. For example, in the experiments of on the HCP-major dataset, by using only 10% of the training set, the training time was reduced 10 times as well. When trained, Classifyber segments bundles in just 3 minutes. The main cost of the computation in both the training and test phases is the preparation of the feature space. For the test phase, almost all of the 3 minutes were spent preparing the target tractogram for the linear classifier (steps (b1) and (b2)), while the actual prediction (step (b3)) only required less than 1 second.

**Figure 10:**
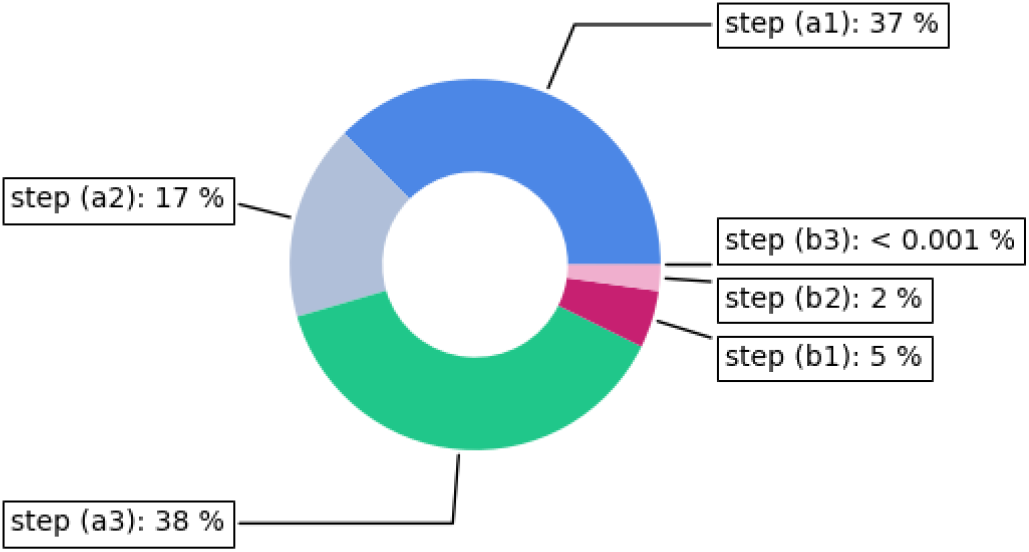
Time to run Classifyber, partitioned into the six steps composing the training phase on 15 subjects and the test phase to predict one IFOF. Total running time: 37 minutes.

In contrast to Classifyber, RecoBundles and LAP do not require training time, because their underlying learning algorithms, i.e. nearest neighbor and linear assignment respectively, are *lazy learning* algorithms that postpone the computation to when the testing/segmentation step is required. In the case of RecoBundles, the segmentation step requires between 0.5 and 3 minutes, on the example discussed above. LAP requires 130 minutes and is thus the slowest of the methods compared.

TractSeg adopts a different approach because it segments 72 bundles in parallel. The training time of TractSeg is vastly larger than all other methods, requiring 7 hours on a GPU. When the bundle of interest is not included in those 72 bundles, or when the training examples differ from the ones used in Wasserthal et al. (2018a), we re-trained TractSeg (called TractSeg-retrained): for example, on the examples of HCP-IFOF, the training phase required approximately 3 hours on GPU, see Table 8. Both TractSeg and TractSeg-retrained required approximately 5 minutes to segment a new bundle.

All computations of all experiments described in this work were executed on the high-performance computing (HPC) cluster provided by Indiana University, allocating 16 cores of Intel Xeon CPU E5-2680 2.50GHz and 32Gb of RAM, a setup equivalent to a powerful personal workstation typically available in research labs and clinics. For TractSeg, we also allocated one NVIDIA GPU RTX 2080Ti.

## 4. Discussion and Conclusions

In this section, we start with general comments on the whole set of experiments with respect to the claims of this work. We then discuss each experiment individually, the effect of segmenting bundles with respect to their fractal dimension (FD), the computational resources needed by each segmentation method, and, before the conclusions, we briefly discuss the issue of reproducing results reported in the literature.

### 4.1. General Comments

At the global level, all the results on the comparison among automatic segmentation methods presented in Section 3.1 indicate one main message: Classifyber clearly outperforms other methods in all cases, by a substantial margin, and segments bundles very accurately. This is observed to occur across different kinds of bundles, tractography techniques, expert-made segmentations, and quality of dMRI data, i.e., research vs clinical quality. The summary results in Figure 8, which report on the y-axes the DSC score for each of the hundreds of individual bundles segmented across all the experiments of Section 3.1, show that Classifyber obtained scores ranging from 0.65 to 0.96, with a mean and standard deviation of 0.85 ± 0.05. This is the highest quality of segmentation among the different automatic segmentation methods by a large or substantial margin, in almost all cases, see the results at the level of individual target bundles reported in Section 3.2.1, 3.2.2, 3.2.3 and 3.2.4 and in Figure 6.

Figure 8 also reports that the results obtained by LAP are consistently superior to those obtained by RecoBundles and TractSeg, at least on the datasets HCP-minor and HCP-IFOF. Moreover, the figure shows that RecoBundles and TractSeg have a large amount of variability in the quality of segmentation across the different experiments: their DSC scores range from 0.07 to 0.90, with means of 0.64±0.14 and 0.71±0.13 respectively. Surprisingly, TractSeg reaches a low (or very low) quality of segmentation on small bundles. We discuss this point in detail below, in Section 4.3, where we discuss the FD of bundles.

### 4.2. Discussion of the comparison across datasets

#### HCP-minor

Figure 6 and Table 3 show that the quality of segmentation obtained by Classifyber is very high (DSC ≥ 0.80) across all kinds of small bundles and distinctively superior to all other methods^6^. This result is of particular importance because minor bundles are notoriously harder to segment due to their size and high variability across subjects (Guevara et al. (2017)).

In the qualitative comparison in Figure 4 we observe that TractSeg-retrained is not very precise in segmenting fine-grained structures of the bundles, in particular their terminal portions. We believe that this is due to an inherent bias of FCNNs, which we discuss in Section 4.3.

#### HCP-IFOF

When segmenting the IFOFs of the HCPIFOF dataset, Classifyber reaches an extremely high quality of segmentation, with DSC around 0.9, as reported in Table 4. RecoBundles and LAP ranked second with DSC around 0.8. TractSeg-retrained, despite being trained on the IFOFs of that dataset, ranked third with DSC around 0.6. Also in this case we believe that this is evidence of an inherent bias of the method, which we discuss in Section 4.3. TractSeg and RecoBundles-atlas ranked last with DSC around 0.5, most probably because the IFOFs in their training set partly differ from the ones in HCP-IFOF, as explained below.

A possible explanation of the poor performances of TractSeg and RecoBundles-atlas is that the anatomical shape of the bundles used as examples differs from the shape of the manually expert-based segmented bundles of the HCP-IFOF dataset. Specifically, the example used by RecoBundles-atlas, i.e. the IFOF of the atlas of Yeh et al. (2018), comes from clustering followed by expert labeling. The examples used by TractSeg come from a semi-automatic refinement of the segmentation provided by TractQuerier (Wassermann et al. (2016)), while the examples in HCP-IFOF are manually segmented by an expert neurosurgeon and follow the definition in Sarubbo et al. (2013) and Hau et al. (2016). These anatomical differences are justified by the fact that the anatomical definition of some white matter bundles, among which the IFOF, is in evolution (Sarubbo et al. (2013); Forkel et al. (2014); Wu et al. (2016a)).

#### HCP-major

Even for the segmentation of major bundles, Classifyber obtained very high quality of segmentation, ranging from DSC = 0.82 for the Right ILF, to DSC = 0.87 for the Right_CST, see Table 5 and Figure 6, outperforming in most of the cases all other methods. Nevertheless, TractSeg reached comparable segmentation quality, with an average DSC ranging from 0.75 to 0.85, even though it used a much larger training set of 84 subjects instead of 15. Due to their size, major bundles are generally easier to segment (Guevara et al. (2017)). On the contrary, RecoBundles-atlas obtained more modest and highly-variable results, with an average DSC ranging from 0.42 to 0.76, although we used the bundle models from Yeh et al. (2018) as suggested by the authors of RecoBundles.

#### Clinical

On the Clinical dataset, i.e. on the white matter of patients with a brain tumor in the same hemisphere as the bundles of interest, Classifyber reached extremely high quality of segmentation, i.e. DSC around 0.9 as reported in Table 6, when the training examples came from the same clinical dataset (see Classifyber-LOSO in the table). When examples partly different from the ones in the Clinical dataset, the DSC dropped accordingly to around 0.8 for Classifyber-IFOF and to 0.7 for Classifyber-major, see for an example Figure 7 (first row). Specifically, in Classifyber-IFOF, the tractography of the training bundles is built on research-quality data instead of clinical-quality and the reconstruction step of the tractography is CSD instead of DTI. In Classifyber-major the differences are even greater: the training data is research-quality and the tractography is probabilistic instead of the deterministic tractography featured in the Clinical dataset. Moreover, in this case, the definition of the IFOF is the classical one provided by TractQuerier (Wassermann et al. (2016)) instead of the more refined from Sarubbo et al. (2013) and Hau et al. (2016) used in the Clinical dataset.

It is well known that training classification algorithms on examples that systematically differ from the examples in the test set substantially reduces the quality of classification. This problem, called *domain adaptation* or *domain shift*, was previously mentioned for bundle segmentation in Wasserthal et al. (2018a) and has no simple solution.

Although in Wasserthal et al. (2018a) they claim that their pre-trained network works properly also on clinical settings, the results of TractSeg on the Clinical dataset are surprisingly low, with DSC around 0.3, as reported in Table 6. These results should be comparable to those of Classifyber-major, which instead reached a DSC around 0.7. We believe that the main reason of this behavior is the low FD of the clinical bundles, which has a strong impact on TractSeg as explained in detail below in Section 4.3.

### 4.3. The fractal dimension of bundles

While conducting hundreds of automatic segmentations with different methods, we noticed that TractSeg had consistent success or consistent failure on specific datasets. TractSeg very accurately segmented the bundles in HCP-major, but obtained only medium or poor results in other datasets, see Figure 6. Figure 8 shows that the segmentation quality reached by TractSeg is deeply affected by a specific geometric property of the voxel mask of the target bundle: its fractal dimension (FD, see Section 2.2.7). Tractseg accurately segmented bundles which are smooth and rounded, i.e., with high FD, while it produced poor segmentations when they are wrinkled and irregular, i.e., with low FD. By Combining the information of Table 7 and the trends in Figure 8, we can indeed expect TractSeg to accurately segment bundles in the HCP-major dataset and to consistently fail in the HCP-IFOF or Clinical datasets.

We believe that this tendency is related to the operations of convolution and max-pooling of the fully convolutional neural networks (FCNNs) within TractSeg. In the domain of computer vision, it has been observed multiple times that FCNNs are biased towards rounded segmentations of objects, which can loose details and fine-grained structure, in particular because of the max-pooling operation, see for example Sabour et al. (2017); Kim et al. (2018); Wei et al. (2019). This problem is inherent in U-net (Ronneberger et al. (2015)), which is at the core of TractSeg.

As an example, consider the experiments related to the segmentation of the IFOFs and Figure 5. The IFOFs in the HCP-IFOF dataset were manually segmented by experts and, according to Table 7, have FD = 1.99 ± 0.06. The IFOFs predicted by TractSeg have FD = 2.3 ± 0.1, and appear substantially more rounded and smoother than the expert-based segmented IFOFs, see Figure 5 (third and last panels). Even re-training TractSeg only on examples of HCP-IFOF did not solve this problem but instead merely mitigated it: the IFOFs predicted by TractSeg-retrained have FD = 2.1 ± 0.1, which is still systematically higher than the expert-based segmentations, confirming the bias, see for example Figure 5 (fourth panel).

Figure 8 shows that LAP is also slightly affected by the FD of bundles, though much less than TractSeg. However, such result might not be entirely reliable because a large portion of the segmentations are missing due to the limitation of LAP to address large tractograms.

In contrast to TractSeg and LAP, both RecoBundles and Classifyber are insensitive to the FD of the voxel masks of bundles, as clearly shown in Figure 8. We speculate that the reason for this is related to the streamline-based nature of such methods and, more specifically, to the fact that they operate via *single* streamline classification. By predicting whether or not each streamline of the tractogram belongs to the target bundle, there is not a specific constraint to produce round/smooth voxel-structures as observed with TractSeg or to jointly consider all target streamlines during the prediction as in LAP.

### 4.4. Size of the training set

As an additional result of this work, we observed that Classifyber requires only a small number of example bundles to obtain high quality of segmentation. In fact, Figure 9 shows that, on the HCP-minor dataset, there is no substantial gain in the quality of segmentation beyond 15 training examples. In the experiments on the Clinical dataset, Classifyber reached an extremely high segmentation quality using only 6 example subjects, with a mean DSC around 0.9, see Table 6.

Both RecoBundles and LAP require a very small number of training subjects: 1 bundle/model for RecoBundles and around 5-10 for LAP, according to Sharmin et al. (2018). On the contrary, TractSeg was trained on 84 subjects. Although in Wasserthal et al. (2018a) there are no clear guidelines on the number of subjects to be used for training, it is well known that deep learning models need a very large training set, which is often not available in clinical settings.

### 4.5. Time required to segment a bundle

Among the methods compared in this work, deciding which one is faster is not straightforward: on the one hand, streamline-based methods like Classifyber, RecoBundles and LAP require the tractogram as input. In our experience and applications, the tractogram is always already available and provided by neurosurgeons/neuroscientists, because they decide the reconstruction and tracking algorithms specifically for their desired task, the available MR scanner and sequence of acquisition. If only raw dMRI data is provided, the time to build the tractogram should be accounted for the total time of the computation. On the other hand, TractSeg uses the GPU and requires a specific pre-processing of dMRI data as input, which needs approximately 30 minutes of computation per subject. Moreover, to obtain the predicted bundle as streamlines, bundle-specific tracking must be computed afterwards (Wasserthal et al. (2018b)).

Overall, if the target tractogram is available, RecoBundles is the fastest segmentation method in our comparison, see Table 8. Alternatively, if pre-trained methods are available, like in the case of TractSeg and Classifyber, TractSeg and Classifyber are also similarly as fast as RecoBundles. LAP is the slowest segmentation method but, if training has to be done, TractSeg ranks last.

### 4.6. Reproducibility

The results on large bundles that we present in Table 5 and Figure 6 accurately reproduce those in Wasserthal et al. (2018a) for what concerns TractSeg and RecoBundles. TractSeg has a distinctively higher quality of segmentation than RecoBundles. However, when considering the dataset HCP-minor, see Figure 6 and Table 3, and the dataset HCP-IFOF, see Table 4, RecoBundles shows comparable quality of segmentation to TractSeg. This result is novel because Wasserthal et al. (2018a) did not consider bundles with low FD.

The better performances of LAP than those of RecoBundles and Tractseg on the dataset HCP-minor and HCP-IFOF are shown in the same tables and figures just mentioned. With respect to RecoBundles, this result is consistent with what was demonstrated in Sharmin et al. (2018), i.e., that LAP outperforms the nearest-neighbor-based segmentation, which is the category in which RecoBundles belongs. With respect to TractSeg, the result of our comparison is novel, because LAP was not included in the extensive comparison presented in Wasserthal et al. (2018a).

The sharing of code and data is becoming standard practice in neuroscience and facilitates both accelerated scientific discovery and reproducibility, see Avesani et al. (2019). For this reason, Classifyber is freely available on the online platform https://brainlife.io both as the full algorithm that implements the training and test phases, and as a pre-trained method ready to segment bundles in the highest quality fashion available.

### 4.7. Conclusions

In this work we present Classifyber, a streamline-based linear classifier that segments white matter bundles from dMRI data and expert-made examples. Classifyber is the first automatic classification segmentation method that exploits both the shape of streamlines, obtained with tractography techniques from dMRI data, and the anatomical information of the bundles, in the form of connectivity patterns and specific ROIs. Classifyber substantially raises the quality of segmentation as compared to the current state-of-the-art methods described in the literature, by a large margin, and more importantly, across very diverse settings. Maintaining a high quality of bundle segmentation regardless of the type of input tractography or the quality of dMRI data is nowadays of paramount importance for a vast number of applications. For example, the practitioner may not be able to anticipate whether the bundle to be segmented will have high or low FD.

As opposed to voxel-based methods, like the one presented in Wasserthal et al. (2018a), we believe that accurate segmentation of bundles from dMRI data must leverage tractography techniques and also include information about streamlines. Streamlines represent a spatial statistic of the dMRI signal that approximates the underlying anatomical connectivity, though it does so with a substantial problem of false positives (Pestilli et al. (2014); Daducci et al. (2015); Maier-Hein et al. (2017); Jeurissen et al. (2019)).

Additionally, Classifyber is fast to train on new datasets/bundles and requires only a small number of examples. This specific feature is of great importance for bundle-specific applications like in pre-surgical planning, because Classifyber can be tailored to the specific task, dMRI data and tractography technique at the cost of a small amount of manual segmentation by expert neuroanatomists.

In future, we plan to test nonlinear classification algorithms in order to investigate potential improvements in the segmentation quality of Classifyber. The current linear model used within Classifyber is indeed a limitation of the proposed method. Nevertheless, linear models are fast and light and, according to the results presented in this work, sufficient to substantially advance the state-of-theart in automatic white matter bundle segmentation.

## Acknowledgements

HCP data were provided by the Human Connectome Project, WU-Minn Consortium (Principal Investigators: David Van Essen and Kamil Ugurbil; 1U54MH091657) funded by the 16 NIH Institutes and Centers that support the NIH Blueprint for Neuroscience Research; and by the McDonnell Center for Systems Neuroscience at Washington University.

F.P. was supported by NSF IIS-1636893, NSF BCS1734853, NSF AOC-1916518, a Microsoft Investigator Fellowship, Microsoft Research Azure Award, Google Cloud Platform, and the Indiana University Areas of Emergent Research initiative “Learning: Brains, Machines, Children.

## Appendix A. Semi-automatic technique to curate the HCP-minor bundle dataset

In this Section, we describe the semi-automatic technique adopted to filter out bundles considered not anatomically plausible.

First, we automatically discarded those subjects which had at least one bundle that deviated more than 2 standard deviations from the mean of the bundle distribution of the number of voxels and number of streamlines of the population across the 192 subjects. After this step, the number of subjects retained was 121. Then, an expert (D.B.) performed visual inspection of each individual bundle to detect anomalies in the segmentations. Bundles were assigned an omnibus score corresponding to their degree of anatomical plausibility. These scores ranged from 1 to 5 such that 1 indicated a rating of *bad*, 2 indicated a rating of *poor*, 3 indicated a rating of *OK*, 4 indicated a rating of *good*, 5 indicated a rating of *great*. Finally, we kept those subjects whose *all* bundles obtained a score of 2 or higher, remaining with a total of 105 subjects.

## Appendix B. Further insights about Methods

### Appendix B.1. Vectorial Representation of a Streamline

Here we provide a comprehensive description of the procedure adopted to transform each streamline into a vector.

Given a streamline *s*, we compute 4 sets of values that, concatenated, create the proposed vectorial representation *v* of the streamline. The first two sets refer to the geometrical aspects of the streamline, typically exploited by streamline-based segmentation methods. The remaining two sets refer to connectivity and anatomical aspects of the bundle of interest respectively, which are the main focus of connectivity-based segmentation methods.

Streamline-based segmentation methods group together streamlines according to some similarity measures, or distances. Typical distances between two streamlines are the *minimum average direct flip* (*d*_MDF_) distance or the *minimum average mean* (*d*_MAM_) distance, which account for the respective positions and shapes of the two streamlines, see Garyfallidis et al. (2015); Olivetti et al. (2017). Based on such concepts, an accurate and easy way to compute a vectorial representation of streamlines has been proposed in Olivetti et al. (2012) and since been used for multiple applications, like clustering, interactive segmentation and fast nearest-neighbor queries, see Olivetti et al. (2013); Porro-Muñoz et al. (2015); Sharmin et al. (2016). The transformation from streamline to vector is built on the general concept of *dissimilarity representation*, initially proposed for pattern recognition problems, see for example the comprehensive Pekalska and Duin (2005). The dissimilarity representation for streamlines described by Olivetti et al. (2012), first requires the user to define a small group of prototypical streamlines of the tractogram^7^, called *landmark* streamlines, *l*_1_, …, *l*_*L*_, that summarises the tractogram and acts as a reference system. Then, given a streamline *s*, the set of its distances from the landmarks is its vectorial representation: *v* = [*d*(*s, l*_1_), …, *d*(*s, l*_*L*_)], where *d* is a streamline distance, like *d*_MDF_ or *d*_MAM_. As shown in Olivetti et al. (2012) and in Porro-Muñoz et al. (2015), a vector *v* or this sort is an accurate vectorial representation of the streamline *s*.

In this work we propose a vectorial representation for streamlines that extends the one originally proposed in Olivetti et al. (2012). The first two sets of values are two dissimilarity representations based on different landmarks: the **first** one uses *L* = 100 landmarks taken *globally* from a whole tractogram, as in Olivetti et al. (2012); the **second** one is bundle-specific and uses *L* = 100 landmarks taken *locally* in the area of bundle of interest. Both the global and local landmarks are chosen in one random subject using the subset farthest first (SFF) policy, which provides a uniform coverage of the area of interest, as suggested in Olivetti et al. (2012). Notice that, since the set of landmarks act as a reference system, they have to be the same for all subjects.

The **third** set of values represents connectivity features and is, again, a dissimilarity representation but now focused on connectivity patterns instead of the shape of the streamline. The idea is that, if a streamline represents the anatomical connection between cortical areas at its endpoints, then two streamlines with neighboring endpoints represent the same pattern of anatomical connectivity and serve the same purpose. The dissimilarity representation of this third set of values is based on a recent streamline distance that we proposed in Bertò et al. (2019), which exploits only the endpoints of the streamline: given two streamlines *s*_*A*_ and *s*_*B*_, whose enpoints are 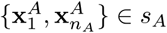 and 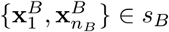, their endpoint distance is simply the mean Euclidean distance of the corresponding endpoints:

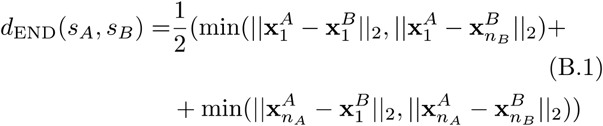

In this work, we propose to use this endpoint distance from the *L* = 100 global landmarks as the the third set of values to describe the *connectivity* pattern of a streamline.

The **fourth** set of values refers to anatomical aspects of the bundle of interest, by means of the ROIs that define that bundle. Often, a bundle is defined by two ROIs, see Wakana et al. (2007); Yeatman et al. (2012); Zhang et al. (2010). In Bertò et al. (2019), we recently proposed a streamline-ROI distance: given a streamline *s* and one ROI represented as a voxel mask ROI = {vox_1_, …, vox_*M*_}, their distance is the minimum among all Euclidean distances between the points of the streamline and the voxels of the ROI:

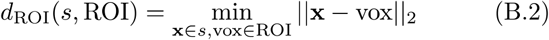

where with vox we indicate the coordinates of the center of the voxel. We use this distance to define the fourth set of values, i.e., the set of distances of the streamline *s* to each of the two ROIs that define the bundle.

In conclusion, given a streamline *s*, we compute 100 values as the dissimilarity representation from *global* landmarks (**set 1**), then 100 values as the dissimilarity representation from *local* landmarks (**set 2**), 100 values as the *endpoint* distance from global landmarks (**set 3**) and 2 values as the Euclidean distance from the 2 ROIs relevant to the bundle of interest (**set 4**)^8^. The vector *v* resulting from concatenating those 302 values is the proposed vectorial representation of the streamline and these 302 variables define the proposed feature space. An illustration of the proposed feature space is given in Figure 2.

### Appendix B.2. Further details about the training and test phases of Classifyber

In this Section, we describe more in details the training and test phases of Classifyber. The *training phase* is composed of three steps:

#### Step (a1) *Bundle superset*

The entire set of streamlines in each tractogram is reduced to a subset of those proximal to the bundle of interest. The main purpose of this reduction is to avoid extremely *imbalanced data*, which decreases the accuracy of classification. Typically, the ratio between the number of streamlines of a bundle (class 1) and all the other streamlines in the tractogram (class 0) is around 1: 500, so extremely imbalanced. A typical simple technique to promote effective training is to remove examples far away from the boundary between the two classes and to get a more even class ratio, which is what we obtained by retaining only the streamlines in the region of the bundle of interest. Such operation is computationally intensive, but we adopted the very fast solution described in Sharmin et al. (2018). Specifically, the bundle superset of an example bundle is computed by considering the neighboring streamlines belonging to the corresponding tractogram retrieved by a *k* nearest neighbors (*k*-NN) procedure applied to each streamline of the bundle. We found *k* = 2000 to be a good compromise between computational cost reduction and size of the resulting superset with respect to the bundle and the tractogram. Usually, with *k* = 2000, the bundle superset, which is a subset of the entire tractogram, is approximately 30 times bigger that the bundle and 20 times smaller than the whole brain tractogram. Efficient *k*-NN queries are possible due to the use of a KDTree, as in Sharmin et al. (2018). Moreover, this extra cost in time is massively outweighed by the 20x gain in time when computing the next steps, i.e. steps (a2) and (a3), see Section 3.2.7 for more details.

#### Step (a2) *Feature extraction*

Each streamline of the superset is then transformed into a vector, as described in Section 2.2.2. To the vector is assigned a class label 1 if it belongs to the bundle, 0 otherwise, see Figure 3 (A), where they are represented in green and red respectively. The entire set of vectors, i.e. the *training set*, is z-scored independently for each feature.

#### Step (a3) *Training*

A binary Logistic Regression classifier is trained, using the stochastic average gradient (SAG) solver (Schmidt et al. (2017)) available in the Python package scikit-learn (Pedregosa et al. (2011)). We use default parameters, except for the number of iterations of the solver, which we increase to 1000 to ensure convergence, as well as the parameter to lessen the negative effects of the residual class imbalance, which we set in all cases to 1:3. These choices are the result of a preliminary investigation on left out data and are kept for all the experiments.

The *test phase* is also composed of three steps:

#### Step (b1) *Bundle superset*

Similarly to step (a1) of the training phase, we reduce the whole target tractogram to a superset of the target bundle. The main reason for this step is to reduce the computational cost of segmenting the target bundle. Obviously, in this case we do not know the target bundle in advance, so the superset is only *expected* to contain the target bundle, with very high probability. In this case, first, a candidate bundle superset is computed as in step (a1) but considering, in the target tractogram, the neighboring streamlines of one of the *example* bundle. This procedure is repeated using 5 of the example bundles. Second, the final bundle superset is obtained as the union of all the candidate bundle supersets. Retrospectively, in all experiments, the superset obtained in this way was approximately 40 times larger than the target bundle and always containing all the streamlines of the target bundle. Step (b2) *Feature extraction*. Each streamline of the bundle superset is embedded into a vector, as described in Section 2.2.2. All the vectors are z-scored feature-by-feature using means and standard deviations obtained in step (a2) of the training phase.

#### Step (b3) *Test*

By exploiting the linear classifier obtained from the training phase in step (a3), each streamline of the superset is predicted to be either part of the bundle (class 1) or not (class 0).

## Appendix C. Further insights about Experiments

### Appendix C.1. TractSeg-retrained metrics on HCP-minor dataset

In Figure C.11, we report the training metrics obtained when training TractSeg-retrained as explained in Section 2.2.5. Red lines represent the value of the loss function obtained across all the epochs (y axis labels on the left side), while green lines represent the f1 score (y axis labels on the right side). The graph shows that we reached convergence in 250 iterations.

### Appendix C.2. Tuning RecoBundles parameters on HCP-minor dataset

Figure C.12 shows a quantitative comparison in terms of mean DSC when using RecoBundles with different configuration of parameters across the HCP-minor dataset. In that specific setting, we found that bundles predicted with default parameters (depicted in light green) obtained, on average, lower DSC scores, with respect to bundles segmented using other two configurations (depicted in yellow and blue). Specifically, the configuration that gave best DSC scores was the one (depicted in blue) that used the minimum average mean distance (*d*_*MAM*_) instead of the minimum average direct flipped distance (*d*_*MDF*_) and that did *not* use the local streamline linear registration (SLR), most probably because the bundles were already coregistered in MNI space. The configuration that used the ‘refine’ option was the one (depicted in light blue) that gave the worst results.

---

**Figure C.11:**
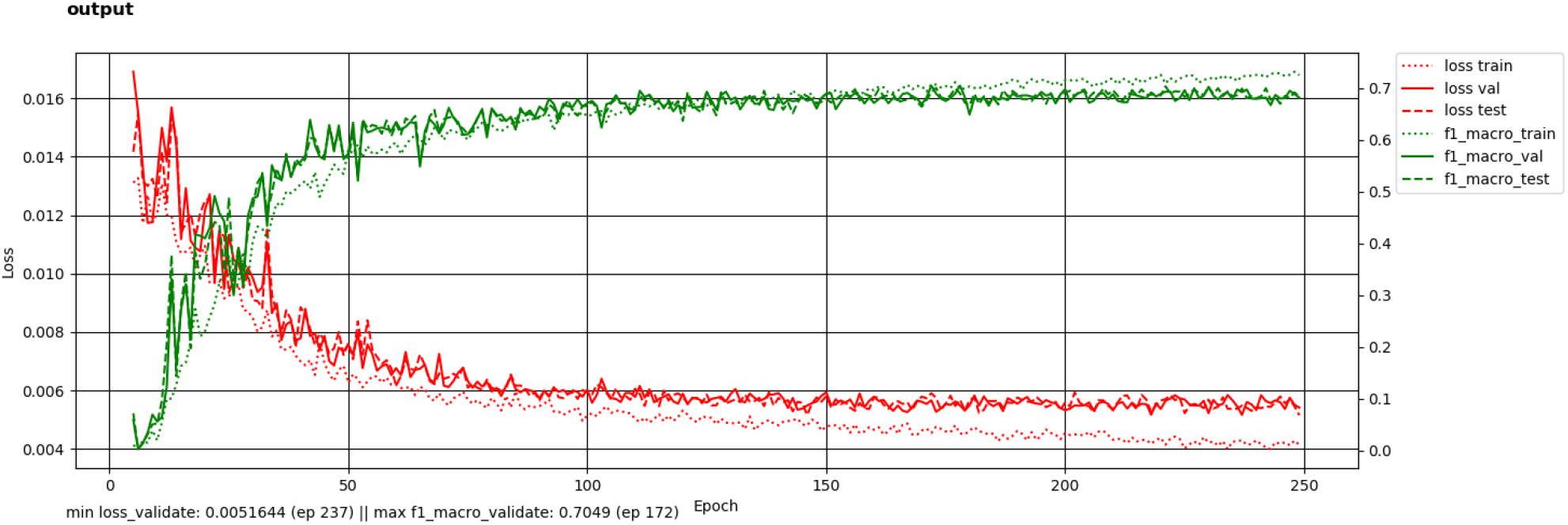
Metrics to train 15 subjects for TractSeg-retrained on the HCP-minor dataset, data augmentation, 250 epochs.

**Figure C.12:**
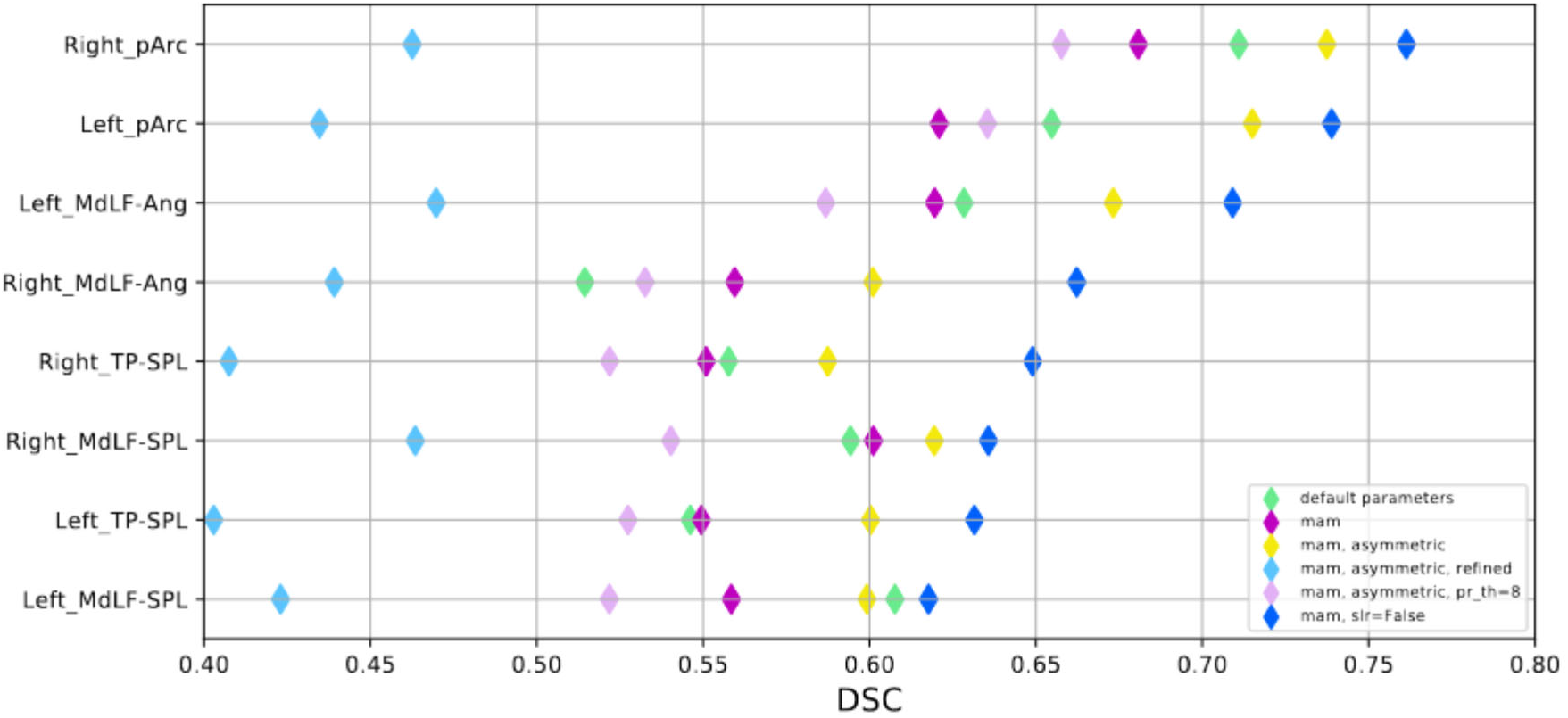
Quantitative comparison of different parameter values for RecoBundles on HCP-minor: mean DSC across 5 target subjects when using RecoBundles with different configuration parameters. Specifically, ‘mam’ means pruning distance=‘mam’ and reduction distance=‘mam’; refined means that was used the refine option; ‘asymmetric’ means slr metric=‘asymmetric’; pr th=8 means pruning threshold=8.

## Footnotes

1 Available at http://brain.labsolver.org.

2 Available at https://figshare.com/articles/FreeSurfer_reconstruction_of_the_MNI152_ICBM2009c_asymmetrical_non-linear_atlas/4223811.

3 Available at https://github.com/vistalab/vistasoft/tree/master/mrDiffusion/templates/MNI_JHU_tracts_ROIs.

4 In some literature, the name *fiber* refers to *axon* and in other literature to *streamline*. Here we refer to the latter for linguistic convenience.

5 Available at http://brain.labsolver.org.

6 The mean improvement in terms of DSC with respect to the second-best method is 0.09.

7 Such streamlines can be just a random subset of the tractogram.

8 Or more than 2 values in case of more than 2 ROIs.

